# Composition efficacy of Unsaturated Arachidonic acid, Diterpenoids, Malvin (C_29_H_35_ClO_17_), and Bergenin to neutralise venom from different venomous snake species

**DOI:** 10.1101/2022.12.07.515639

**Authors:** Lujaina N. H. Al-Tobi, Juma.Z.K Albusaidi, Ali.A Ajabri, Mohammed A. Idris, Sidgi S. A A. Hasson

## Abstract

Snakebite envenomation is a serious problem in tropical and subtropical countries. Antivenom is the only treatment used to treat snake envenomation, however it is unable to neutralise local haemorrhage. Therefore, this study’s aim is to evaluate the efficacy of *P. dulce* leaf extract to neutralise local haemorrhage induced by three clinically important snake species, *B. jararaca, C. atrox* and *E. carinatus*. Moreover, to determine the active components which are responsible for this activity. The plant leaves were extracted using different solvents, however, only E/e extract showed the best neutralizing capacity. The increasing doses, DF-1:2; 1:4, of E/e extract allowed better neutralizing ability s.c. In contrast, the oral/ i.p. acute toxicity test revealed that the optimal doses for the administration of E/e were 1 and 8 mg/kg. In addition to that, E/e was tested for its anti-lathality of LD_50_ using *B. jararaca* venom (1.1mg/kg) i.p., where the higher doses of 16 and 24 mg/kg killed 75% of BALB/C mice. Consequently, the different components of E/e extract were isolated with HPLC. The different components were grouped and tested to uncover the active ones. The results revealed that only three fractions were active, Frc11, Frc13, and Frc14. The active fractions showed a disparity in neutralizing the individual venoms, however, the best neutralising capacity was scored for Frc11. When the same fractions were pooled together, they showed a complete neutralizing ability against individual venoms as well as the pooled venoms. That was confirmed with the anti-gelatinase activity test, where pooled fraction inhibited the SVMP enzyme which is responsible for gelatinase activity. The phytochemical characterisation showed that the active fractions consist mainly of secondary metabolites such as tannins and polyphenols. MALDI-TOF MS confirmed the presence of secondary metabolites in the active fractions. The same fractions were tested for their anti-lethal activity using the pooled venoms (LD_100_), the results were statistically not significant, as all mice died including the positive controls. Nevertheless, the active fractions showed a noticeable increasing in survival time period especially Frc13 with an average survival time of 37 minutes. The positive control, IAV, scored the longest survival period with a gap of 11 minutes from Frc13.

## 1. Introduction

Snake envenomation is a serious health problem in rural tropical and subtropical countries (Gutiérrez, et al., 2006; Kasturiratne et al., 2008; Menaldo, et al., 2015; Moreira, et al., 2016), affecting mainly young (Kasturiratne, et al., 2008), poor farmers (Calvete, et al., 2009; Gutiérrez, et al., 2006). Populations in these regions experience high morbidity and mortality due to several factors, such as poor access to health services, which are often suboptimal (Kasturiratne, et al., 2008) and, in some instances, limitation and unavailability of antivenom (Chippaux, 2008; Kasturiratne, et al., 2008), which is the only specific treatment (Stock, et al., 2007), due to market failure (Chippaux, 1998; 2008). Many of survivals suffer from long-term physical disability due to local tissue necrosis and psychological seqeulae (Kasturiratne, et al., 2008; Calvete, et al., 2009). This environmental and occupational health problem has not received the adequate attention from national and international health authorities (Kasturiratne, et al., 2008), therefore it has been added to the list of the neglected tropical diseases (NTD) (WHO, 2007; Hifumi et al., 2015).

Antivenom is the only specific treatment for snakebite envenomation (Arora & Choudhary, 2016; Barreto et al., 2017; Chippaux et al., 1998; Gutiérrez et al., 2006; Kasturiratne et al., 2008). Antivenom has the ability to reverse systemic complications of snakebite envenomation. However, it cannot reverse local haemorrhage as well as other clinical complications such a local and systemic tissue necrosis; such as pre-synaptic neurotoxicity (nerve terminal destruction) (Kuruppu et al., 2008), myotoxicity or renal injury (Isbister et al., 2008). The failure of antivenom in treating such complications is reasoned by: antivenom inefficacy, irreversible venom-mediated effects, its inability of to reach venom target, rapid venom onset as well as mismatch of venom and its pharmacokinetics. The main reason attributed to that i.e. the drawbacks is the small molecular weight of potent toxin molecules, which allow them to migrate faster deep in the tissues. Hence, the advantage of the fragmented antibodies rely on their small molecular weight and its ability to migrate deep in tissue, but cannot treat such devastating injuries stated above i.e. pre-synaptic neurotoxicity and myotoxicity. The other second and crucial concept is that, the highly toxic molecules in the venom are that which have small molecular weight, thus weakly immunogenic compared with non-toxic molecules. Therefore, the highest titer of antibodies is directed against non-toxic molecules. Thirdly, some of the snake venoms are highly dangerous and have rapid venom onset. Black mamba for example is one of the dangerous snakes and their venom makes collapse within 45 minutes. Finally, it is also related to the relation between the mismatch of venom and antivenom pharmacokinetics. Whole immunoglobulins (IgG) have higher molecular weight, which increase their half-life. However, fractioned immunoglobulins have lower molecular weight and their elimination rate even faster. And this factor leads to recurrence of the venom toxins, especially those with longer half -life i.e. intravascular toxins. In addition to the previously mentioned disadvantages, whole antibodies antivenoms (IgG) were previously thought to cause adverse reactions due to the presence of Fc portion (Gutiérrez et al., 2011). Interestingly, they found that both whole and fragmented antibodies can cause adverse reactions. And the variation of the severity depends on the physiochemical features of the antivenom. Antivenoms with high physiochemical quality are clear of non-IgG contaminating proteins as well as protein aggregates. Such antivenoms have low incidence of adverse reactions, caused by complement system. On the other hand, there is low tolerability experienced with antivenoms of poor physiochemical features i. e. turbidity, high content of protein aggregates or non-IgG contaminants (Gutiérrez et al., 2011). Moreover, there is a limitation and shortage in the production of antivenom (Chippaux, 2008; Kasturiratne et al., 2008; Maduwage et al., 2016), due to market failure (Chippaux, 2008). Furthermore, snakebite incidences occur mostly in rural tropic (Chippaux, 2008; Kasturiratne et al., 2008), where antivenoms are neither available nor affordable.

The use of complementary and alternative medicine has become popular throughout the world. It represents a considerable part of the health care provided worldwide. The use of traditional medicine has reached 80% and 65% in low and high income countries, respectively. Due to the previously mentioned drawbacks of the antivenom, finding an alternative agent becomes a necessity. The main objective of our study, therefore, is to evaluate the efficacy of *Pithecellobium dulce (P. dulce)* leaf extract to cross react with venoms of different snake species and neutralise their local haemorrhage effects.

## 2. Materials & method

### 2.1 Plant material

The leaves were obtained from different regions since the plant is widely distributed in Oman. The leaves were picked carefully to make sure that do not contain any type of infection. Then, they were washed to remove any dust or contamination, followed by shadow drying in a hot room for 3 days at 50°C. The third day, the leaves were collected, crushed, and weighed. The total dry weight was 78.27 g divided equally between 3 solvents using sterile (500 ml) bottles.

### 2.2 Extraction of plant material

For the preparation of *P. dulce* leaves extract, three major different solvents were used which are, ethanol (C_2_H_6_O, absolute ≥99.8%), methanol (CH_3_OH; 97.9%) and autoclaved distilled water. The dry weight of the crushed powdered leaves used for each solvent was 26.1g. The mixture was incubated at room temperature (23°C) with continuous shaking for 7 days. On the seventh day, the extracts were sieved into three a separate beaker. Each of the sieved solutions went into a further filtration using a funnel and Whatman filter paper. Then, each of these extracts was dried under reduced pressure to the minimal volume possible using rotary evaporator. Next, each of the extracts was separated into three portions and each was re-extracted with the same solvents to end up with nine extracts.

### 2.3 Source of venom

Three venoms from clinically important snake species were used and selected which include; *C. atrox* 100mg (V7000-100MG - USA), *B. jararaca* 10mg (V5625-10MG - Brazil), and *E. carinatus* 10mg (V8250-10MG - France) were studied. The venoms were purchased in a powder form (lyophilized) from Sigma. Prior to use gradual amounts of autoclaved distilled water was used to liquidize the venom powder, where *C. atrox* venom was diluted to get a ratio of 2mg/ml. While each of *B. jararaca* and *E. carinatus* were prepared to get 1mg/ml.

### 2.4 Ethics

The ethics committee approval was obtained from the Ethical Committee, Sultan Qaboos University (SQU). The instructions of the ethical committee as well as the international standardization on using experimental animals were strictly followed. This including that the animals were handled carefully and exposed to a minimal pain and stress.

### 2.5 Image J

A software program used to measure the haemorrhagic area following the instructions of Tang et al., (2010).

### 2.6 Minimum haemorrhagic dose

To determine the MHD of the venom, *in vitro* test was done. An amounts of 0.3, 0.6, 0.8 and 1μl were taken from *B. jararaca* (1mg/ml). While the amounts taken from the remaining species were 6, 8, 12 and 16μl. Each of these volumes was top up with autoclaved distilled water to get a final volume of 200μl. The tubes were then incubated in a water bath at 37°C for 30 minutes. The venom was injected subcutaneously on hind limbs, with different concentration.

### 2.7 Verification of the active extract

This step was done to determine the most efficient extract that gives 100% neutralization efficacy. The different solvents extracts were prepared by weighing and dissolving the dry crude material (100mg) with autoclaved distilled water (900μl) to get a ratio of (1:10), in separate labeled tubes. An amount of 192μl was taken from the previously prepared extracts and placed in separate Eppendorf tubes. An amount of 16μg from *C. atrox* venom (2mg/ml) was added to each of the different mixtures and pre-incubated in a hot water bath with the venom at 37°C for an hour. Eighteen BALB/C mice were used to be injected subcautansely on hind limbs, each limb with 200μl, Sixteen hours later, animals were sacrificed and their skins were dissected to observe haemorrhage.

### 2.8 Acute toxicity test

#### 2.8.1 Oral route

The optimal dose of *P. dulce* leaves extract that show no toxicity effects, will be determined using *in vivo* test. Twenty BALB/C mice of both sexes (25g) were grouped into four groups (5 mice/group) and dosed orally with increasing doses (1, 8, 16 and 20 mg/kg) of extracted sample. The control group of 5 mice will be used in parallel and receive an equivalent volume of distilled water. Animals were watched for signs of acute toxicity and death over 24h.

#### 2.8.2 Intraperitoneal route

A similar experiment will be performed using the intraperitoneal (i.p.) route. For each extract sample a total of 20 mice of both sexes were distributed randomly into four groups and treated i.p. with increasing doses (1, 8, 16 and 20mg/kg) of the extract. Animals were observed over 24h for signs of acute toxicity and death. A control group was treated with an equivalent volume of distilled water following the same route. The survival time of the mice will be recorded and analysed using Log-rank test.

### 2.9 Determination of the effective dose (ED) subcutaneously in vivo

*In vivo* test analysis was performed to determine the ED of E/e extract. In this experiment, *B. jararaca* venom was used. And the MHD of this venom was determined to be 0.3μl (0.3μg). Before starting the experiments, an amount of 100mg of E/e extract was dissolved with 900μl of diluents (1:10). Once the stock solution was prepared, different volumes were taken to prepare different concentrations. Each of these amounts was dissolved in diluents to get a total volume of 200μl (Table 3.3). Because, the total amount injected in one hind limb was 200μl. Therefore, the amounts mentioned previously were doubled for each mouse. Once the venom was added to all concentrations, they were incubated in water bath at 37°C for an hour. BALB/C mice were injected subcutaneously on hind limbs. After 16 hours, animals were sacrificed and their skins were dissected to observe the neutralization efficacy.

### 2.10 High performance liquid chromatography (HPLC)

HPLC was used using C18 column (BUCHI, 40-63μm) to fraction the whole extract (2:1 ratio) into its components. HPLC is a separation technique and its principle depends on injecting a small amount, of 5ml of the liquid extract in each turn, into a HPLC column. This column consists of small particles made of silica in which polar components are attached to. HPLC uses two different solvents, which are normally polar such as water and acetonitrile. The solvents washed out the polar compounds, which in return will be detected by UV light (280nm) to determine the absorbance of each fraction and shown as peaks in the system chart. Once the absorbance of each fraction has been detected, they are loaded into test tubes and are known as the stationary phase. The fractions, which their absorbance is beyond the threshold, are considered as a waste and will be loaded into the waste beaker. The total fractions collected were 173 fractions.

#### 2.10.1 Screening of fractions

The collected 173 fractions were grouped into 9 groups, each consists of 20 fractions, except for the last group (13 fractions), in order to minimize the number of mice sacrificed. Nine Eppendorf tubes were labeled alphabetically (A, B, C, D, E, F, G, H, I, J, K) i.e., group “A”, fractions range (1-20), “B” (21-40), “C” (41-60) and so on. Each of the previous groups were preincubated with the MHD of *B. jararaca* venom at 37°C for an hour. BALB/C mice were injected subcutaneously on hind limbs. Sixteen hours later, animals were sacrificed and their skins were dissected to observe the neutralization efficacy.

### 2.11 Characterization of the active fractions

#### 2.11.1 Sodium dodecyl sulfate-poly acrylamide gel electrophoresis (SDS-PAGE)

SDS-Page was performed to indicate the presence of proteins in the E/e extract as well as the active fractions. The size of the protein ladder (PL) used was up to 42 kDa.

#### 2.11.2 Phytochemical test

##### Tannin content analysis

The presence of tannins in E/e extract and active fraction(s), was determined using tannin test. In this test, ferric chloride solution (FeCl_3_) was used. Development of blue-black colour indicates the presence of hydrolysable tannins. While, the formation of greenish colour represent a combination of hydrolysable and condensed tannins (Pithayanukul et al., 2005).

##### Alkaloid content analysis

Determination of alkaloids was also performed using Mayer’s reagent. Mayer’s reagent is prepared by dissolving 1.36g of mercuric chloride and 5g of potassium iodide into 100mL distilled water and mixed well. An adequate amount of Mayer’ s reagent was added to E/e extract as well as the active fractions. The presence of alkaloids is indicated by formation of cream colour precipitate (Gunatilaka et al., 1980).

##### Flavonoids content analysis

To determine the presence of flavonoids in E/e extract as well as the active fraction(s), sodium hydroxide solution (NaOH) was used. An amount of 1ml of NaOH solution was prepared (1:10) as a stock solution. A diluted amount of the stock solution was added to the extract and the active fractions. A formation of condensed yellow colour indicates the presence of flavonoids. And the intensity of the colour decreases when a diluted acid is presented. (Hossain et al., 2013).

#### 2.11.3 Matrix-assisted laser desorption/ionization-time of flight mass spectrometry (MALDI-TOF MS)

MALDI-TOF MS was used to determine and identify the biomolecules present in the active fraction(s). The MALDI-TOF MS is a method found in the beginning of 1980s. This method influenced greatly in the increasing the applicability of mass pectrophotometry (MS). The principle depends on ionizing the protein sample. And this performed by mixing the sample with an energy-absorbent, organic compound called matrix. The matrix crystallizes on drying, and so do the sample, which is trapped within the matrix. The trapped sample is ionized automatedly by pointing a laser beam into the sample. Laser beam ionizing process results in the formation of protonated ions from analytes in the sample. Those protonated ions travel at a fixed potential, where they are separated according to their mass-to-charge ratio (m/z). Then, charged analytes are detected and measured with time of flight analyzer (TOF). Based on TOF results, a biomolecules mass fingerprint (BMF), characteristic spectrum, is generated for analytes in sample (Singhal et al., 2015).

#### 2.11.4 Gelatic zymography

Gelatin zymography is a technique used to detect the activity of gelatinase enzymes e.g. matrix metalloproteinases following the method of Hasson et al., 2004. Snake venoms are rich with hydrolases enzymes, therefore determination of the active fraction(s) activity to neutralize snake venom can be detected using this method. Active gelatinases will degrade gelatin in gel and appears as clear bands against the dark stained background. On the other hand, neutralising activity of the active component(s) should stop the hydrolysing enzymes and appears as dark. The *B. jararaca* venom used in this experiment was prepared with cold 1x PBS (phosphate-buffered saline, pH 7.2). The concentration of the venom which was prepared was 5mg/1ml (5: 1 ratio). (Hasson et al., 2004).

#### 2.11.5 Anti-lethal activity

Median lethal dose (LD_50_) was used to determine the anti-lathal activity of E/e extract of *P. dulce* leaves. Different concentrations of E/e extract, 2, 4, 8, 12, 16 and 24 mg/kg, were preincubated with LD_50_ of *B. jararaca* venom (1.1mg/kg). BALB/C mice (25g) were injected i.p. with the mixture and monitored for 24 hours. The active fraction(s) were tested against the complete lethal dose (LD_100_) of pooled venoms. They were preincubated with the pooled venoms. BALB/C mice were injected i.p. with the mixture and were monitored for 24 hours.

#### 2.11.6 Statistical analysis

The statistical analysis was performed using IBM SPSS statistics 23. The data were obtained from Image J by calculating the area of the haemorrhage. Statistical analysis of the inhibition of the haemorrhage as well as survival rate in mice compared with the control groups was performed using analysis of variance, Manwetny test, to determine the significance (*p*<0.05) of the data obtained in all experiments.

## 3. Results and analysis

### 3.1 Results criteria

For the concept of understanding, the analysis of the hemorrhage was based on the criteria that established accordingly, Table 3.1, and in line with the international measures. The haemorrhage was examined with a naked eye and followed by a confirmation methodology where. The haemorrhage images were assessed by using a software program, Image J. Image J software calculates the surface area of the haemorrhage (mm^2^) to support and confirm the intensity of the haemorrhage by observing.

**Table 3.1:**
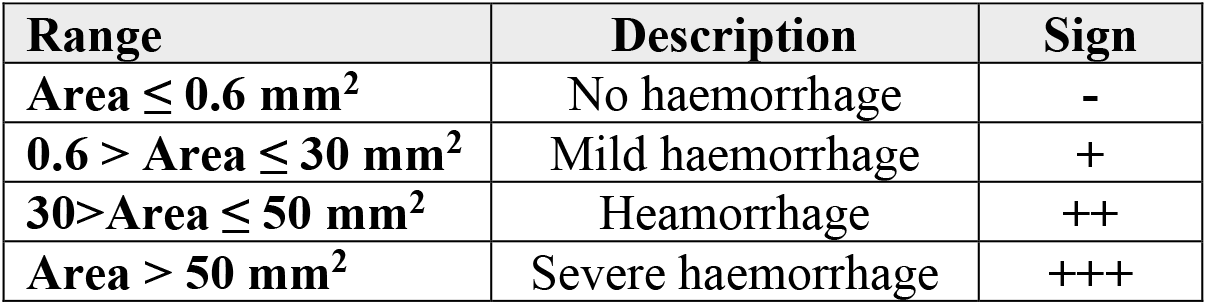
Differentiation criteria between mild, to severe haemorrhage.

| Range | Description | Sign |
| --- | --- | --- |
| $\text{Area} \leq 0.6 \text{ mm}^2$ | No haemorrhage | - |
| $0.6 > \text{Area} \leq 30 \text{ mm}^2$ | Mild haemorrhage | + |
| $30 > \text{Area} \leq 50 \text{ mm}^2$ | Heamorrhage | ++ |
| $\text{Area} > 50 \text{ mm}^2$ | Severe haemorrhage | +++ |

### 3.2 Evaluation of the minimum haemorrhagic dose (MHD)

The first investigation was carried out to explore the required dose, for each individual venom that will give MHD i.e., 50mm^2^ to 280mm^2^. The results were found to be as follows. The MHD of *B. jararaca* was determined to be 0.3μg. While, each of *C. atrox* and *E. carinatus* venoms were determined to be 16 μg and 8 μg, respectively. However, the choice of the haemorrhagic area range depends on the potency of the snake venom.

### 3.3 Verification of the effective extract

The *P. dulce* plant leaves was extracted using different solvents as Ethanolic/methanolic (E/m), Ethanolic/water (E/w), and methanolic/water (M/w) extracts. The efficiency of each of the extracts to neutralise local haemorrhage was determined by injecting BALB/C mice subcutaneously, detailed in chapter 3. The results as illustrated in Figure 3.1 and 3.2 showed that the Ethanolic/ethanolic (E/e) extract was found to be significant in neutralising the venom compared with the other groups. All alcoholic extracts showed neutralisation of local haemorrhage. The same thing was relatively seen with methanolic extracts. However, two of the aqueous extract, W/w, W/m, were unable to overcome the haemorrhage. The only extract which showed remarkable inhibition of local haemorrhage was E/e extract. E/e extract showed protection against local haemorrhage induced by snake venom in both groups in comparison with the other extracts. The results were further analysed using the Image J software as shown in Figure 3.2.

**Figure 3. 1:**
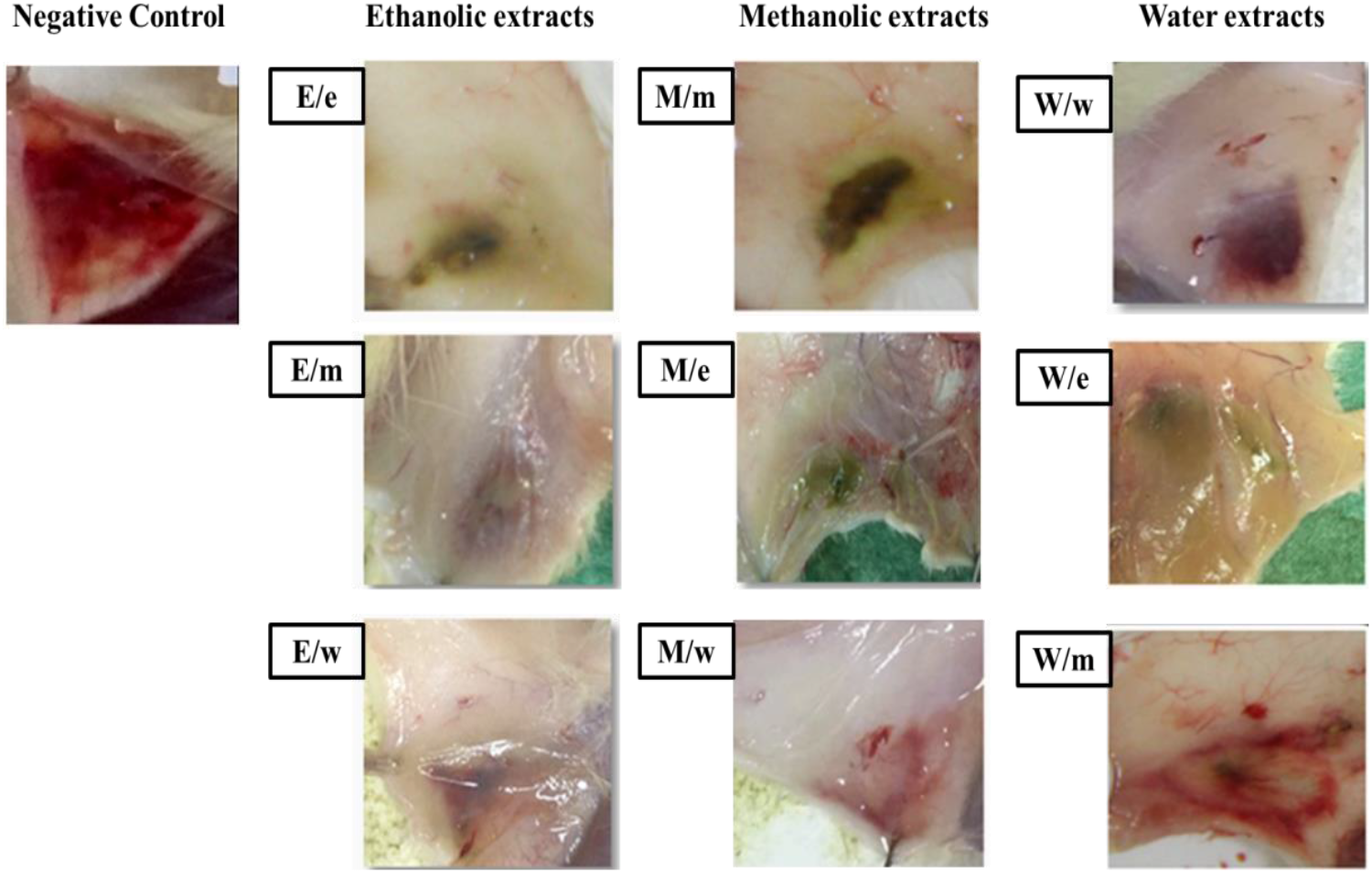
Verification of the active extract. E/e: Ethanolic/ethanolic; E/m: Ethanolic/methanolic; E/w: Ethanolic/water; M/m: Methanolic/methanolic; M/e: Methanolic/ethanolic; M/w: Methanolic/water; W/w: Water/water; W/e: Water/ethanolic; W/m: Water/methanolic.

**Figure 3. 2:**
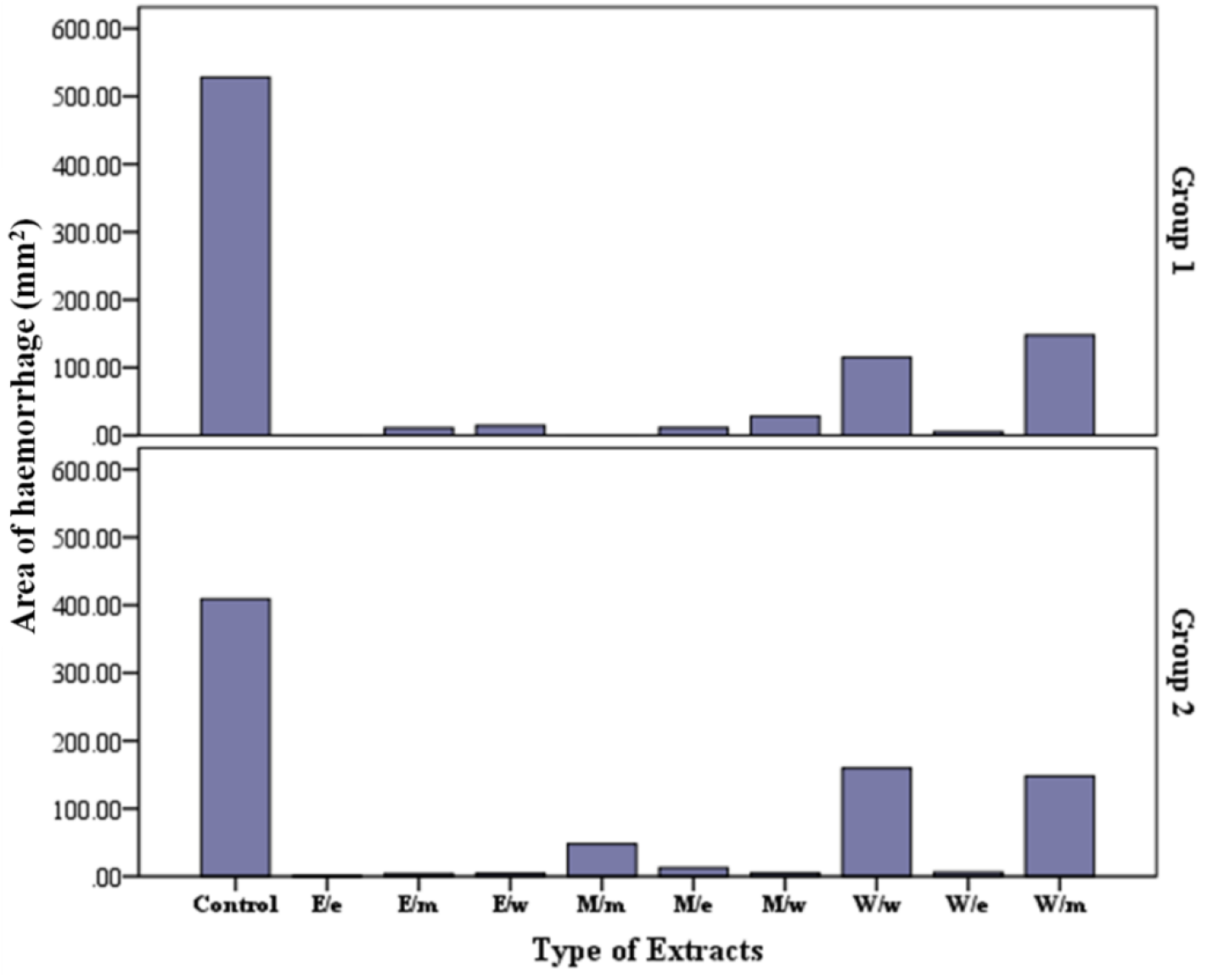
Screening of different solvent extracts. E/e: Ethanolic/ethanolic; E/m: Ethanolic/methanolic; E/w: Ethanolic/water; M/m: Methanolic/methanolic; M/e: Methanolic/ethanolic; M/w: Methanolic/water; W/w: Water/water; W/e: Water/ethanolic; W/m: Water/methanolic.

### 3.4 Determination of the optimal dose for oral/i.p. administration

The E/e extract was tested for its acute toxicity as well as to determine the optimal dose for oral/i.p. administration by introducing an increasing doses of E/e extract. It was noticed that higher doses cause a rapid depolarisation and death. While lower doses of 1 and 8 mg/kg showed no toxicity to animals. However, whether this toxicity is associated with the small animals (i.e. mice and rats) only, or may affect larger animals including humans is still obscure.

### 3.5 Determination of the effective dose (ED) of E/e

The determination of the ED was done by introducing different concentrations (1:20, 1:10, 1:4 and 1:2) of E/e in BALB/C mice subcutaneously. All the concentrations used were found to be significant in nuetralising the haemorhage induced by the snake venom. Although the lower concentrations of 1:20 and 1:10 showed partial inhibition of the haemorrhage as illustrated in Figure 3.3. However, the 100% neutralistion efficacy was a concentration dependent manner by which the dilution factors (DFs) 1:4 (Figure 3.3d) and 1:2 (Figure 3.3e) showed a complete (100%) neutralisation activity of the local haemorhage induced by the *B. jararaca* venom. However, at lower concentration, the area of haemorrhage declined sharply to reach 6.5mm^2^ and 1.7mm^2^ in both groups, respectively. Therefore, the concentration is proportional with the neutralisation activity subcutaneous rout. The results were further processed using the Image J software as shown by Figure 3.4.

**Figure 3. 3:**
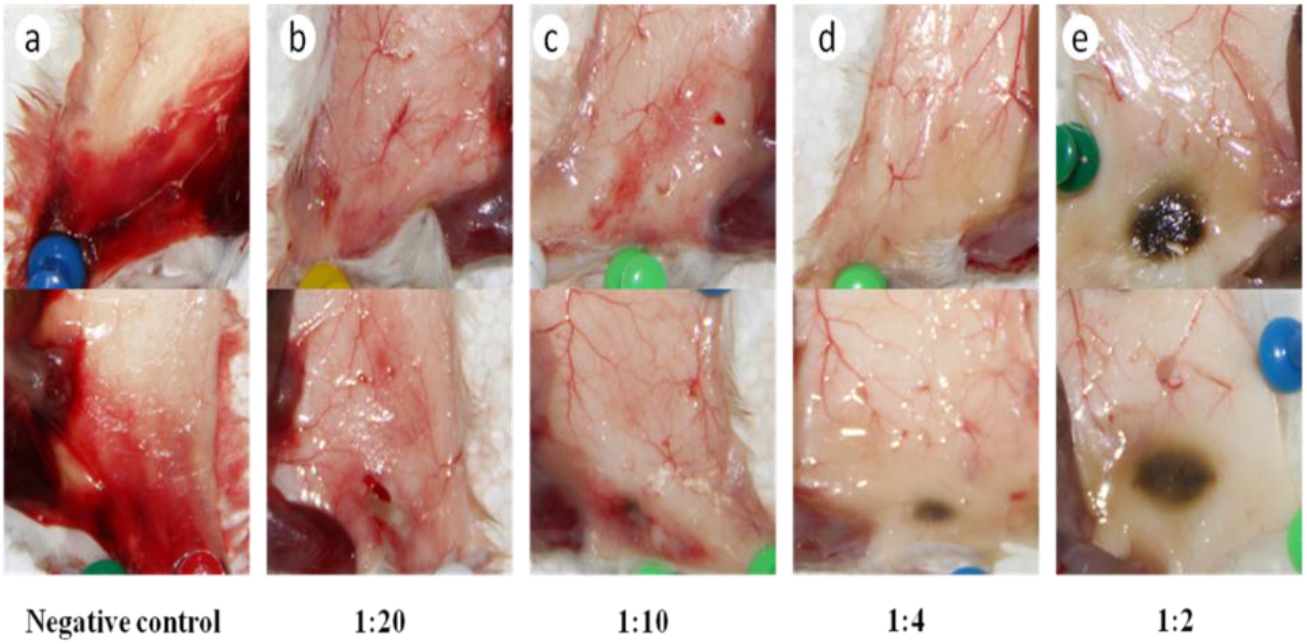
Histological sections show efficacy of different DFs of the E/e crude extract to neutralise the local haemorrhage induced by *B. jararaca*. a) represents the negative control which contains venom only. While, (b-e) are the preincubated mixture of different concentrations of E/e with the venom.

**Figure 3. 4:**
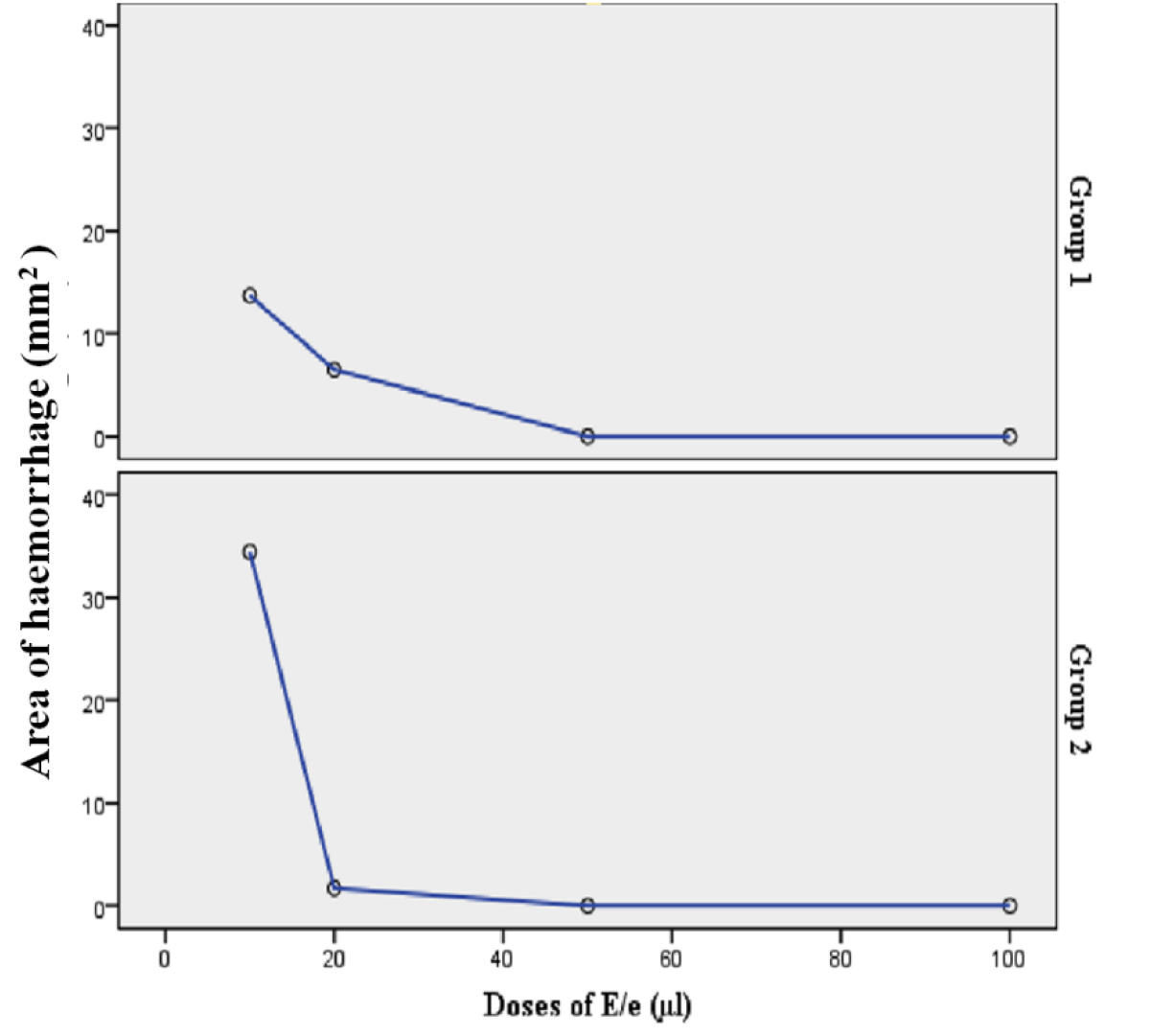
Linear relationship of the different DFs of E/e with their neutralisation efficacy.

**Figure 3.5:**
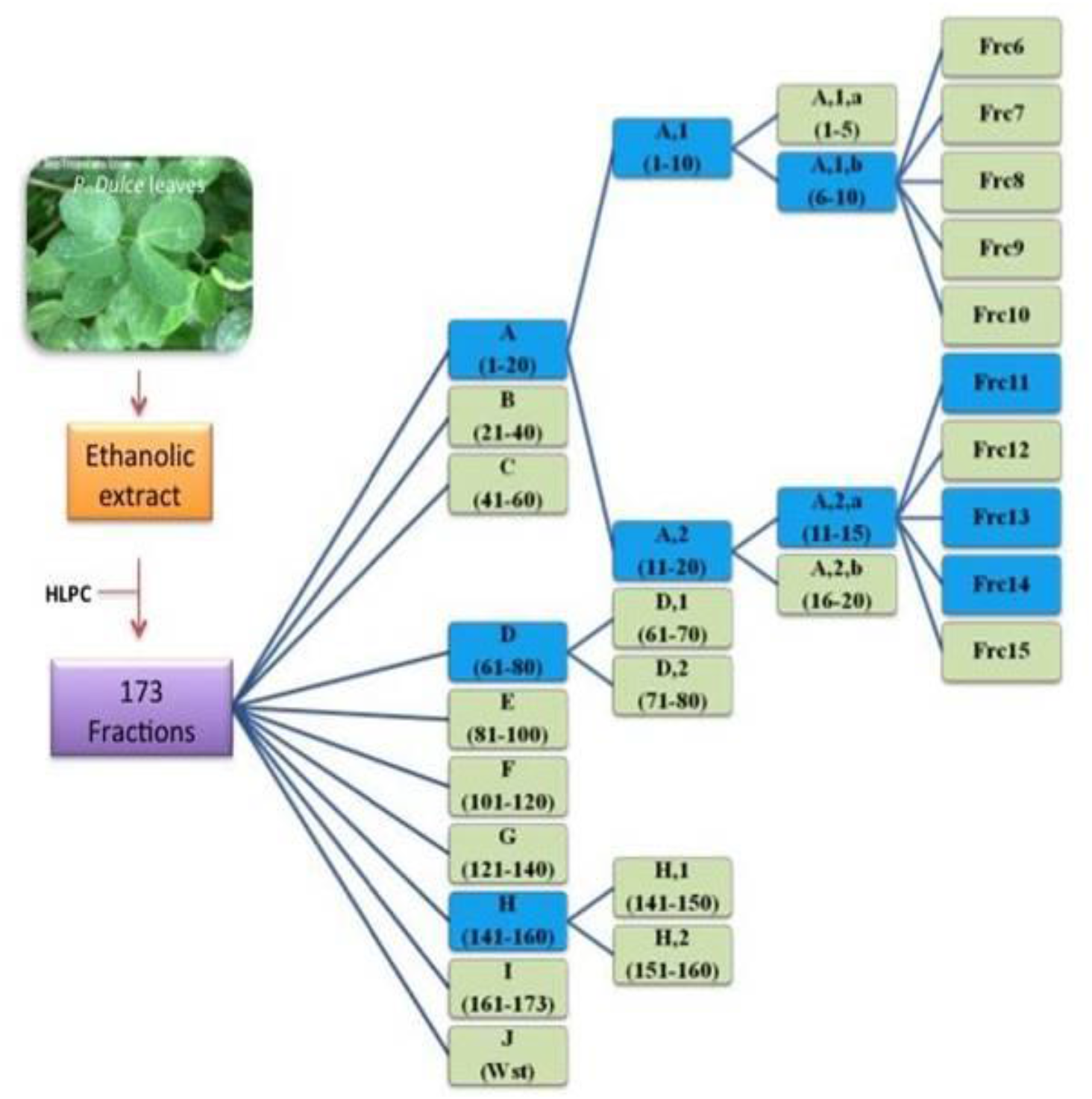
Grouping of the obtained fractions and evaluation of their efficacy.

### 3.6 Fractionation of the E/e extract using High performance liquid chromatography (HPLC

HPLC was used using C18 column (BUCHI, 40-63μm) to fraction the whole extract into its components. The fractions obtained were 173 as illustrated. The analysis of the fractions were grouped and analysed into 9 groups as illustrated in the schematic diagram figure 3.5. Each of the 9 groups consists of 20 fractions. All fractions were tested against *B. jararaca* primarily. Out of the 9 groups only 3 groups, A (1-20), D (61-80) and H (141-160), were found to be effective to neutralise the haemorhage effects induced by the *B. jararaca* venom. Interestingly when each of the effective 3 groups used to investigate the neutralisation efficacy against pooled venom, only one group i.e., group A was found to have the potential candidate(s) fraction(s).

This group, i.e., group A which consists of 20 fractions, was further sub-grouped into two groups as A1 and A2 each of which consist of 10 fractions. When examining both subgroups the results were found to be significant and interesting both groups A1 and A2 neutralised *B. jararaca* venom. Groups A1 and A2 were further subdivided into A1a, A1b, A2a and A2b, 5 fractions in each sub-group, and the desire fraction(s) was found both groups of A1b and A2a. Subgrouping was carried out for this subgroups accordingly aiming to find the potential fraction individually. The results of this further subgrouping revealed that when pooling three individual fractions, 11, 13 and 14 of the main group A2 stated above, were found to have the potential to cross react and neutralise the pooled venoms *B. jararaca*, *C. atrox* and *E. carinatus*. However, when each fraction being examined against individual venom the results were found to be surprising, as each fraction neutralised the individual and pooled venoms differently as showed in Figure 4.6,a & b. Each of the 11, 13, 14 fractions and pooled fractions were able to neutralise each of *B. jararaca* venom. Although fraction 11 and pooled fractions have the ability to neutralise *E. carinatus* venom, fractions 13 and 14 were found to be partially effective (Figure 3.6a). On the other hand, all the individual fractions showed no neutralisation against *C. atrox* and pooled venoms. Surprisingly the pooled fractions showed a complete neutralisation of *C. atrox* as well as pooled venom.

**Figure 3.6:**
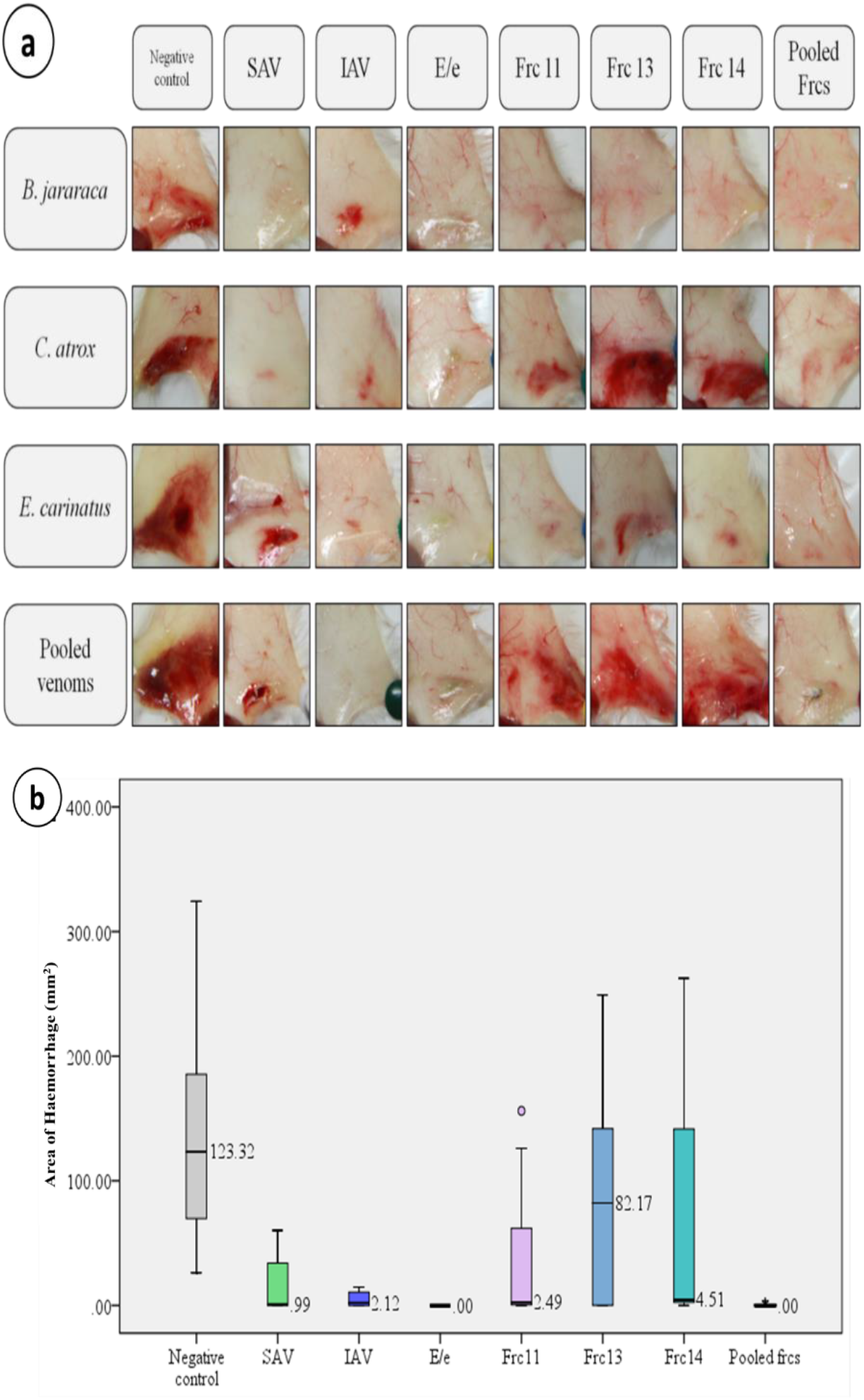
evaluation the efficacy of the active fractions 11, 13 and 14 against B. jararaca, C. atrox, E. carinatus as well as pooled venoms. (a) Shows the histological samples mice dissected skins. (b) neutralisation activity of the fractions 11, 13, 14 as well as the pooled fractions to neutralise different venoms in comparison with the positive and negative controls. Box plot shows the area of haemorrhage. The graph clearly shows that fraction 11 has the best neutralisation activity compared with the other two fractions. When the fractions are pooled together, they showed a complete netralisation of the venoms. *SAV: Saudi antivenom; IAV: Indian antivenom; E/e: Ethanolic/ethanolic; Frc/s: Fraction/s*.

For further confirmation an egg embryo neutralization assay was performed in order to insure the neutralisation activity of the active fractions 11, 13 and 14. Each of these fractions were examined as individual and pooled fraction against *B. jararaca* venom.

Fractions were also analysed for their absorbance intensity in order to calculate their concentrations. The absorbance of the three fraction 11, 13 and were found to be 1.5, 1.18 and 1, respectively. The concentrations therefore determined to be; Frc11 (0.026mg/mL), Frc13 (0.021mg/mL) and 14 (0.018mg/mL). Frc11 was found to be highly concentrated hence the absorption was found to be higher than the other two fractions.

### 3.7 Characterization of the active fractions

#### 3.7.1 Sodium dodecyl sulfate poly acrylamide gel electrophoresis (SDS-Page)

SDS-Page was performed to indicate the presence of proteins in the E/e extract as well as the active fractions. The size of the protein ladder (PL) used was up to 42 kDa. Nothing worth mentioning except for the blue clog that was noticed in E/e and Frc14 lanes may be related to the high concentrations of low molecular weight proteins that had been denatured due to the exposure to high temperature during the process of extraction. On the other hand, samples of Frc11 and Frc 13 showed an empty lanes, which indicates the absence of protein in any of the samples. It can be concluded that it’s unlikely the outcome of the results of neutralisation was due to the presence of proteins.

#### 3.7.2 Phytochemical screening

A phytochemical screening profile was done for the crude E/e extract as well as for the active fractions, 11, 13, 14 to evaluate the presence of tannins, alkaloids and flavonoids. There are several types of tannins such as hydrolysable and condensed tannins. The presence of hydrolysable tannin for example will form a bluish black colour. While Formation of brownish green colour indicates the presence of condensed tannin. Whereas, the presence of both types of tannins will develops olive green colour. The rsults of this assessment was found to be that the E/e extract contains both hydrolysable and condensed tannins (HCT). On the other hand, Frc11 contains condensed tannin (CT) only, while Frc13 and Frc14 both consist of hydrolysable tannins (HT) as illustrated in (Table 3.2). In addition to that using Mayer’s test, all the samples showed negative results in regards of the presence of alkaloids (Table 3.2). Moreover, all samples were tested for the presence of flavonoids using sodium hydroxide. All the samples were positive for flavonoids (Table 3.2).

**Table 3.2:**
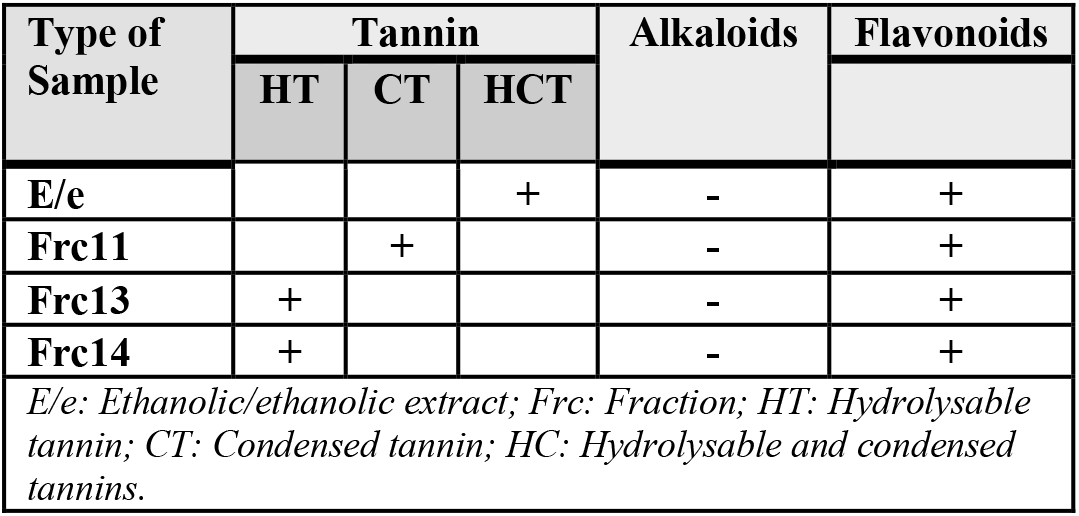
Determination of the presence of phytochemicals in E/e and the active fractions.

| Type of Sample | Tannin |  |  | Alkaloids | Flavonoids |
| --- | --- | --- | --- | --- | --- |
|  | HT | CT | HCT |  |  |
| **E/e** |  |  | + | - | + |
| **Frc11** |  | + |  | - | + |
| **Frc13** | + |  |  | - | + |
| **Frc14** | + |  |  | - | + |
| *E/e: Ethanolic/ethanolic extract; Frc: Fraction; HT: Hydrolysable tannin; CT: Condensed tannin; HC: Hydrolysable and condensed tannins.* |  |  |  |  |  |

**Table 3.3:**
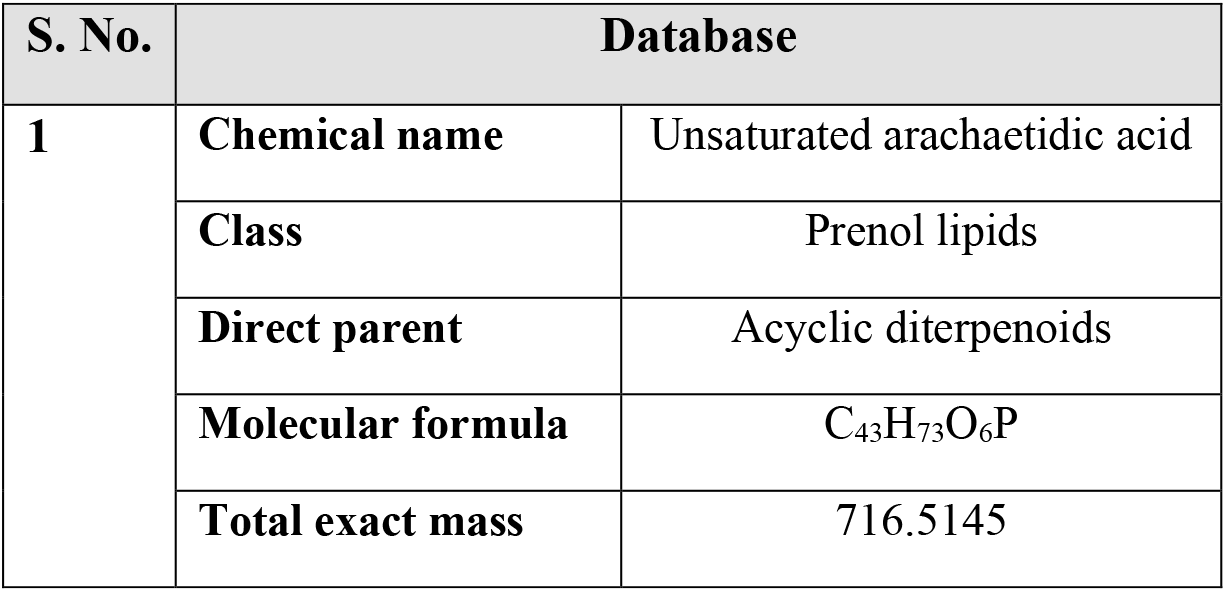
The different components of the peaks of Frc11.

**Table 3.4:**
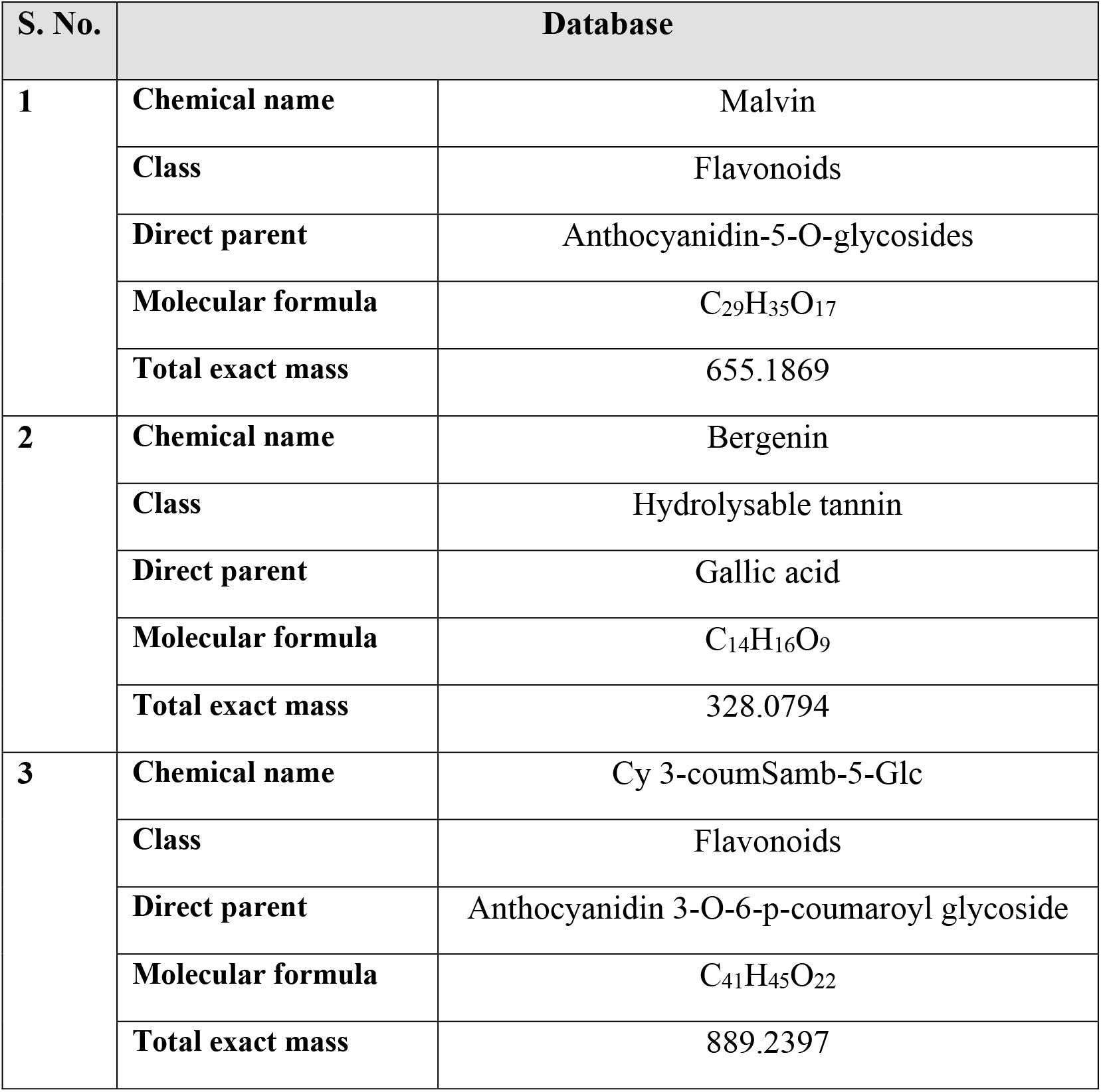
The components of the peaks presented by both Frc13 and Frc14.

#### 3.7.3 Matrix assisted laser desorption ionisation-time of flight-molecular weight analysis (MALDI-TOF)

MALDI-TOF is a technique used to identify and determine the mass-to-charge ratio (m/z) of unknown biomolecules. The samples, Frc11, Frc13, Frc14, were mixed with appropriate matrix to crystallise the sample. The samples were struck with laser beam to evaporate the ions of the sample. Those evaporated protonated ions travel along tube and separate according to their m/z. Ions with lower molecular weight (MW) travel faster and are detected first by the detector. Therefore, mass-to-charge ratio is proportional to the MW.

MALDI-TOF was done in the central analytical and applied research unit (CAARU) in SQU. The analysis was performed using α-cyano-4-hydroxycinnamic acid (HCCA) matrix and Sinapinic acid (SA) matrix. HCCA and SA are used to prepare samples matrices. The spectra were obtained in the range between 600 Da to 200 kDa. This step was done to indicate the nature of the biomolecules present in the active fractions.

After MALDI-TOF was performed, several peaks were obtained. Frc11, showed two peaks of 806.135 and 972.976m/z. The X-axis represents the m/z, the mass of the material divided by its ion charge. And in MALDI-TOF, the m/z is considered as a mass; because the ion charge is always one either it was positive or negative. Therefore, a website was used to determine the molecular formula and the type of the material from the m/z value, www.massbank.jp. While Y-axis represents the resolution and it depends on the size of the sample. The larger the sample, the lower resolution we got. The following shows the beaks obtained for each fraction.

### 3.8 Gelatin zymography

Gelatin zymogram gel is a method used to detect active gelatinase enzymes i.e. matrix metalloproteinases (MMP). The gel is incorporated with gelatin to detect the active gelatinase within the samples. Such samples tend to degrade the gelatin and appear as light bands against the dark stained background.

Snake venoms are rich with MMPs enzymes which influence greatly the local necrosis. This step was done to determine the ability of the samples to neutralise venom MMP enzymes. From the results shown in figure 3.7, it be clearly seen that the clear lane along *B. jararaca* venom is a sign for MMP enzyme activity. The same thing was noticed along the lanes of Frc11, Frc13 and Frc14. However, the lane of “pooled fractions” strongly inhibited gelatinase enzyme activity in the same manner of the positive controls SAV and IAV.

**Figure 3.7:**
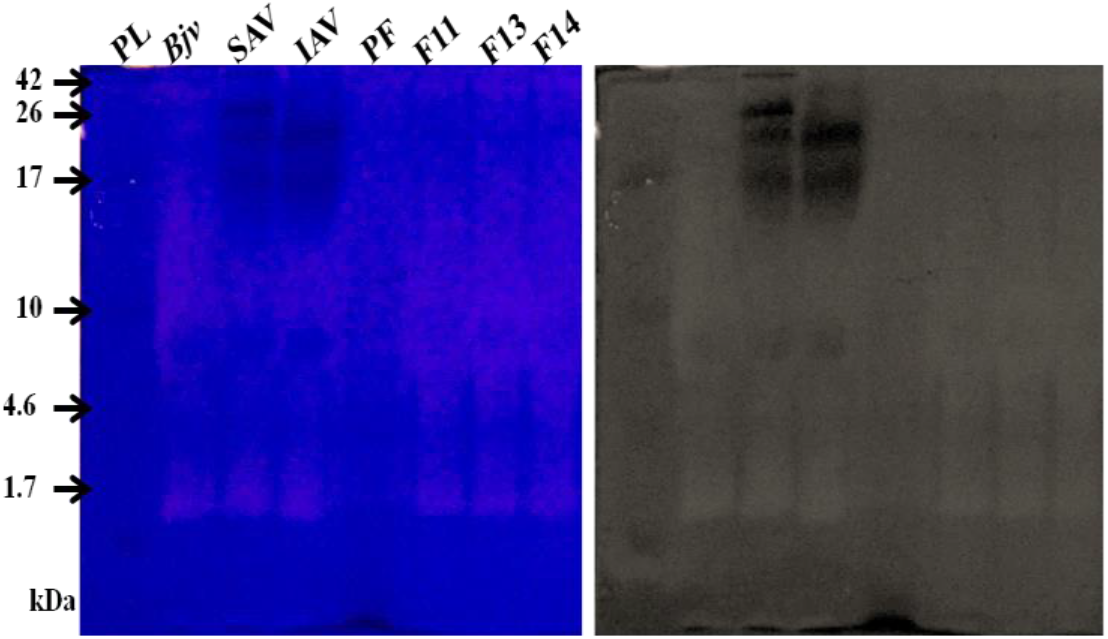
Determination of anti-gelatinase activity of the active fractions as well as the pooled fractions against *B. jararaca* venom. PL: protein ladder; Bjv: B. jararaca venom; SAV: Saudi antivenom; IAV: Indian antivenom; PFrcs: Pooled fractions; Frc: Fraction

### 3.8 Anti-lethal activity

The E/e extract and the active fractions were evaluated for their anti-lethal activity i.e., death prevention. The median lethal dose (LD_50_) of *B. jararaca* was used to evaluate the efficacy of different concentrations of E/e using *in vivo* assay. BALB/C mice being injected intraperitoneally (ip). Prior the experiment illustrated in Table 4.10, a LD_50_ of *B. jararaca* (1.1 mg/kg) was worked out and found to be 27.5μg for a mouse weight of 25g. The results obtained were quite interesting. It was noticed that the survival rate found to be decreased at higher concentrations of E/e extract. Each of the doses 50 and 200μg showed a significant survival rate *(p<0.05)* compared with the other concentrations (Table 3.5). Surprisingly at 100 and 300μg, 50% of mice died. While on the highest doses, 5000 and 9000μg, most of mice died (75%).

**Table 3.5:**
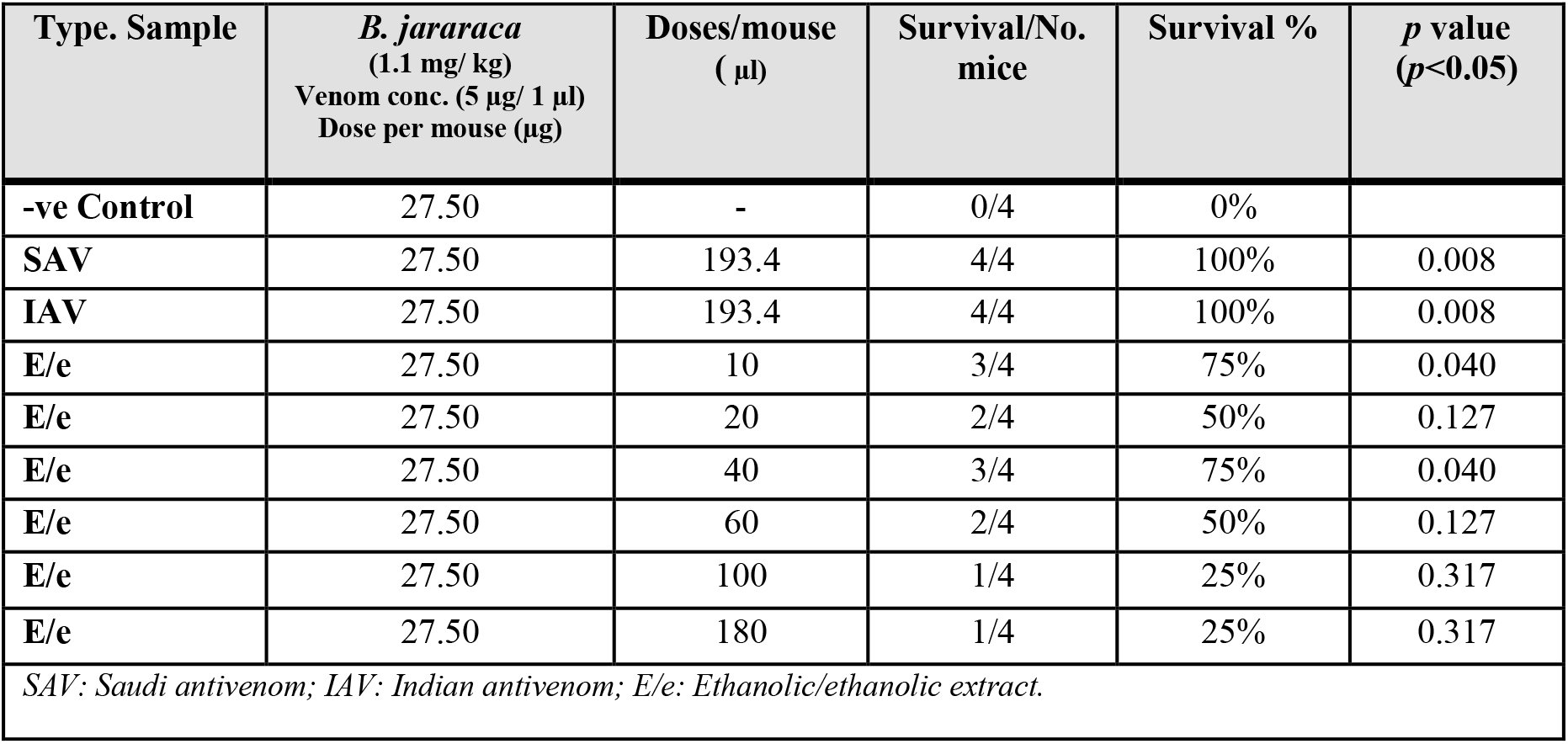
The anti-median lethal activity of the different concentrations of E/e.

The complete lethal dose (LD_100_) was evaluated by using pooled venoms from the three species used. The venom was pooled with 30μg of *B. jararaca* (5mg/1ml), 100μg of *C. atrox* (2mg/1ml) and 50μg of *E. carinatus* (1mg/1ml). E/e extract and the active fractions were tested for their efficacy to neutralise LD_100_ of the pooled venoms. The active fractions, the E/e extract, positive controls (SAV and IAV) along with negative control were evaluated for their anti-lethal activity. All samples used including the positive controls failed to prevent mice death. The average survival time of negative control was 18 minutes, followed by Frc11 and the commercial SAV with an average of 24 and 29:28 minutes. Frc13 survived the longest time compared with the other two fractions with an average of 37 minutes. The same time was relatively scored for E/e extract and Frc14 with 35 and 33 minutes, respectively. In contrast, mice which received the commercial IAV survived for an average of 48 minutes (Table 3.6).

**Table 3.6:**
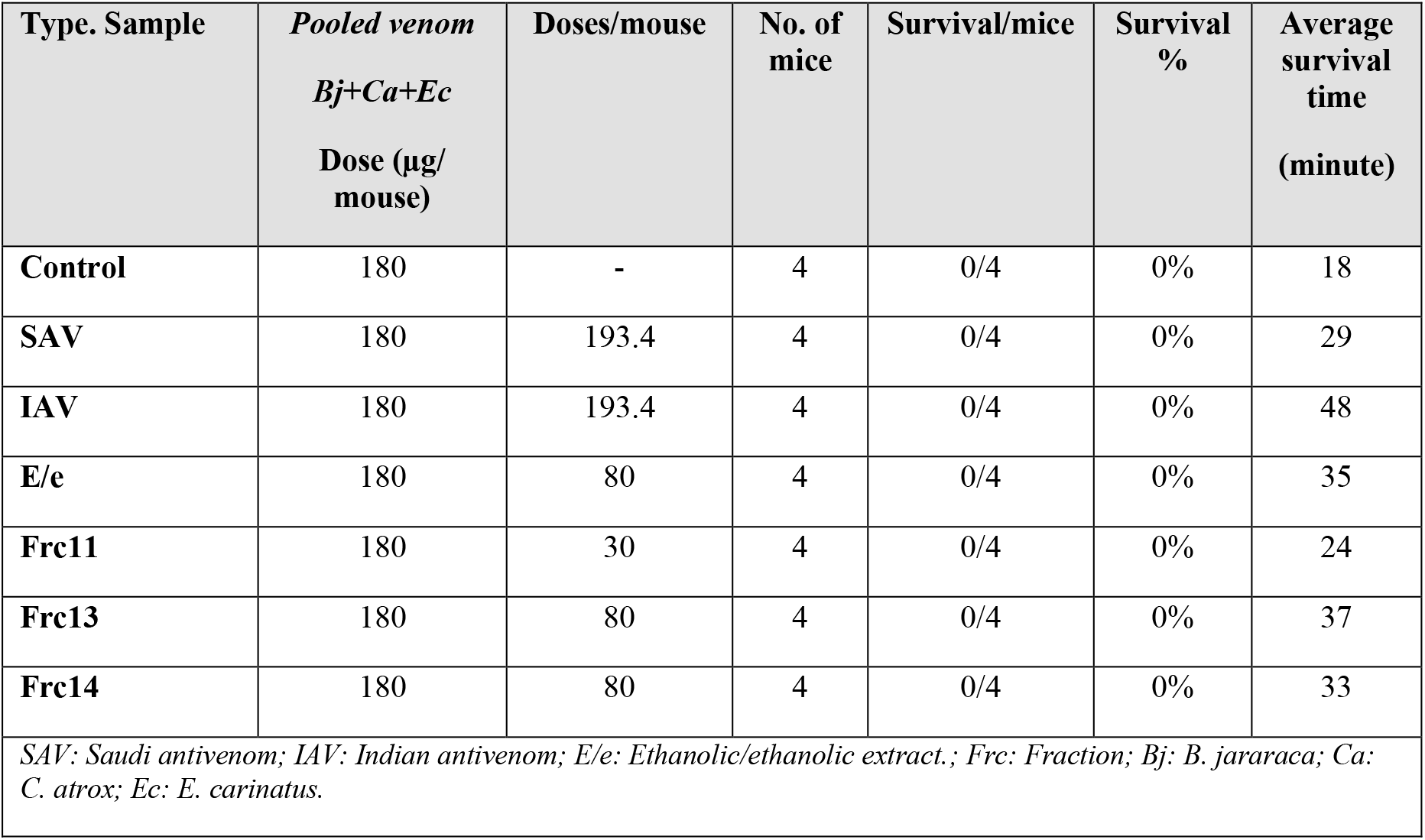
The anti-total lethal activity of the E/e and the active fractions.

## 4. Discussion

Snakebite envenoming leads to serious systemic and local complications (Tanwar et al., 2017). Administration of whole immunoglobulin antivenom intravenously is the only treatment used to prevent systemic pathology. However, such antivenom has limited effectiveness against local envenoming, that results to severe pain, swelling, blistering, haemorrhage and necrosis (Cardoso et al., 1993), which usually cause long-term disability. The inability of IgG antivenom to treat such injuries is attributed to the rapid onset of venom toxins as well as their low molecular weight compared with IgG antivenom which enable them to cross blood barriers faster (Ismail et al., 1998; León et al., 1999; Gutiérrez et al., 2003). Although fragmented antivenoms Fab and F(ab)2 are of smaller size, but they unable to treat local envenoming, whether they were administered by intravenous or intramuscular routes (Theakston, 1997). Therefore, researchers and scientists have worked to develop a treatment for local envenoming focused on the application of natural (Samy et al., 2012) or synthetic inhibitors (Lipps et al., 1996) to overcome snake venom powerful molecules. A recent antivenom known as Específico Pessoa has been isolated from a plant known locally as cabeça de negro. This antivenom has been produced in Brazil and widely used in North and Northeast of the country to treat snakes and scorpions envenomations (Moura et al., 2014). This will reflect the main objective of this research work “Herbal Medicine” as a source of alternative and effective treatment of such problems. Because natural products from medicinal plants may give a new source of medication, there are many research groups that are now engaged in medicinal plants research not only for the discovery for new drugs, but possibly for discovering compounds with novel mechanisms of action that can stimulate new fields of research.

Some of these plants are *Quercus infectoria* (Gupta & Peshin, 2012; Lim, 2012; Shabbir et al., 2014), *Azima tetracantha* (Bordon et al., 2015; Janardhan et al., 2014), *Allium cepa* L, *Annona senegalensis, Bidens pilosa* L, *Conyza sumatrensis, Corchurus trilocularis L,Furestia Africana, Grewia sp.* (Owuor & Kisangau, 2006), *Bombacopsis glabra* (Bhattacharjee & Bhattacharyya, 2013; Mendes et al., 2013), *Piper longum* L (Shenoy et al., 2014) and *Euphorbia hirta* (Gopi et al., 2016; Gopi et al., 2015). Their role of action has been well explained in the literature to be due to several phytochemicals including polyphenols and tannins. In the case of *P. dulce* is known for their pharmacological activities. This in agreement with other studies which also admit the presence of other phytochemicals such as: alkaloids, saponins, steroids (Manna et al., 2011), glycosides (Nagmoti et al., 2012), anthroquinones, terpenoids and sterol (Kalavani et al., 2016). Furthermore, polysaccharides (Peethi and Saral, 2016) as well as saturated and unsaturated fatty acids have been reported by Kalavani et al., 2016.

In this study *P. dulce* leaf is the part of the plant that has been investigated for their anti-haemorrhagic activity against the MHD of different snake venoms. The MHD of the snake venoms used was determined to be 0.3μg, 16μg and 8μg of the *B. jararaca, C. atrox* and *E. carinatus*, respectively. The surface area of haemorrhage (mm^2^) was determined using Image J software to access and confirm haemorrhage analysis. Such doses resulted of an area of haemorrhage ranges from 50-280mm^2^. The variability of the haemorrhage zone is due to unstabilty distruibution and spread of enzymes action within the venom or the size of the animal and their level of tolerability. Hence the MHD was chosen to be the does at which the area of haemorrhage ranges from 5-28 cm^2^.

The *P. dulce* leaf was extracted using different solvents include: EthOH, MeOH and water. Generally, the ethanolic extract showed a remarkable neutralising activity. The severity of haemorrhage in methanolic extracts was mild (+) in contrast with the M/e and M/w extracts. The same thing was relatively noticed when using a mixture of E/W or E/M extracts, only pure ethanolic extract showed a significant anti-snake venom activity. The reason can be attributed to those alcoholic solvents, especially ethanol, can readily dissolve phenols and polyphenols. And those molecules are known for their antioxidant activity (Megala and Geetha, 2010). Their mechanism of action is explained by their ability to chelate metal ions, which are strictly tangled in the production of free radical. The hydrophopic benzenoid rings and hydrogen-bonding potential of the phenolic hydroxyl groups increase the ability of phenols to interact with proteins. This explains their ability as an antioxidants and their ability to inhibit some enzymes by chelating metal ions e.g. zinc ion in the SVMPs, which is one of the potent molecules exist in the snake venom. The main classes of phenolics are Phenolic acids and Flavonoids (Pereira et al., 2009).

The E/e extract was tested for acute toxicity and to obtain the optical dose for oral/i.p. route. BALA/C mice experienced CNS depression at the higher doses of 16 and 24 mg/kg, where most of animals died. While at the lower doses of 1 and 8 mg/kg showed no side effects for the animals. Therefore, the optimal dose for oral/i.p. administration is from 1-8 mg/kg. A study was performed by Sugumaran et al., (2008), compared the acute toxicity of *P. dulce* leaves caused by alcoholic extract and benzamine extract. Alcoholic show more toxicity than any other solvents used. And reasoned the toxicity to increasing of γ-Aminobutyric acid (GABA) in the brain.

The E/e extract has showed complete inhibition of toxins-inducing haemorrhage in *B. jararaca*. This inhibition was clearly observed when high dose - DFs of 1:4 and 1:2 of E/e extract was used. The first sign of envenomation by *B. jararaca* is rapture of blood vessels, causing extravasation of blood contents resulting in local and/or systemic haemorrhage, caused by haemorrhagins e.g. SVMP. And this leads to an inflammatory response. Inflammation is likely occur in response to the haemorrhage or by the PLA2 induction of tumer necrosis factor α (TNF-α). The activity of E/e extract as an anti-haemorrhagic increased in response to the increase in concentration of E/e extract intradermally. This was confirmed by the results obtained when the DF was used at 1:20, where an area of haemorrhage zone was found to be between 13mm^2^ to 34mm^2^. Whereas, lowering the DF to 1:10 resulted of haemorhagic zone between 1mm^2^ to 6mm^2^.

The phytochemical content tests of the crude extract showed that the extract consist of polyphenols. The results showed that the E/e extract were found to contain flavonoids, hydrolysable and condensed tannins. Pithayanukul et al., 2005 reported that the mechanism of action could be due to the binding of polyphenols with venom proteins. The author also claimed that the presence of both tannins enhances the antivenom activity of the extract. Therefore, the high concentrated polyphenols possess a better neutralising activity of the extract.

The anti-lethal activity of the crude leaf extract was tested against the LD_50_. Different doses, 2mg/kg, 4mg/kg, 8mg/kg, 12mg/kg, 16mg/kg, 24mg/kg, of E/e solution were used to assess the LD_50_ activity against the *B. jararaca* venom (1.1mg/kg) via i.p route. The results were quite interesting as a complete inhibition was observed at lower doses of E/e extract and where a significant protection of mice from death was achieved. This was totally opposite and in a different concept of that obtained previously as stated above. However, the higher doses were found to be highly toxic for the mice as its lead to kill the mice instantly. The doses of 2mg/kg and 8mg/kg showed the highest survival percentage with 75%. While the doses of 4mg/kg and 12mg/kg resulted in 50% deaths. The highest doses, 16mg/kg and 24mg/kg, were found to be highly toxic as it killed 75% of mice. The only possible explanation for this may be that *P. dulce* cause toxicity at high concentration to BALB/C mice with further complication with the snake venoms. A clear observation in declining of mice activity [as start with gradual to a complete depolarisation followed by death] was found to be associated with higher doses of E/e extract. Sugumaran et al., 2008 performed a study about the toxic effects of *P. dulce* leaves to albino mice. Their results showed that the cause of CNS depression, was due to the increasing the concentration of GABA in the brain (Sugumaran et al., 2008). In addition to that, leaves are known in general by having many toxic properties. They have abortificient, spermicidal, adulticidal and larvicidal activity. And they are rich in toxic elements such as Arsenic, Cadmium and Lead (Khanzada et al., 2013). Therefore, isolation of the active component would be anticipated and essential to avoid any side effect or clinical complications.

The majority of the current studies that carried out in discovering the antivenom activity of medicinal plants stopped at the point of the crude extract activity. The novelty of present study was to explore further, investigate and isolate the active component(s) within the crude extract. The separation of the E/e crude extract was achieved using HPLC. A total of 173 fractions were obtained, grouped and tested. The active fractions were all from the same subdivision, A2a, Frc11, Frc13 and Frc14.

The isolated active components where further characterized for SDS-PAGE was performed to determine whether they are proteins or not. However, the results obtained were negative. Crude E/e extract that showed a blue blog mass in parallel with fraction 14 which could be due to high concentrations of the samples. MALDI TOF analysis confirmed the absence of proteins in any of the samples isolated. Therefore, it’s unlikely the neutralising activity of the active fractions and E/e extract was due to the presence of proteins.

Fraction samples were further analysed using the MALDI-TOF MS to identify the biomolecules present in the active fractions. Two matrices were used, HCCA and SA and the spectra used was from 600Da to 200kDa. Two beaks were obtained ranged from 600Da to 2000Da from Frc11, revealing that Frc11 consists of two components, a lipid (Unsaturated arachaetidic acid) and a derivative of diterpenoids. Frc13 result in ten beaks, however two only were identified due to the system data base available. It consists mainly of flavonoids (Malvin, Cy 3-coumSamb-5-Glc) and hydrolysable tannin (Bergenin). While Frc14 graph end up with eight beaks, only two were identified due to the system data base available. It consists of the same components as Frc13.

Moreover, the fraction samples were further charecterised for their phytochemical analysis tests to base on the presence of flavonoids and tannins. Frc11 has condensed tannins, while each of Frc13 and 14 has hydrolysable tannins. All fractions contain flavonoids. This explains partially of the antivenom activity of those fractions.

Subsequently, these different fractions were tested individually and in combination against three different snake venoms. There was a disparity in the neutralising activity of the fractions. For example, Frc11 showed the best neutralisation activity to neutralise *B. jararaca, E. carinatus* venoms and partially inhibited *C. atrox* and pooled venom. The present of Acyclic diterpenoids molecule in Frc11 may be responsible to have this role of action. Diterpenoids are known for their anti-inflammatory properties. A report performed by Januario et al., 2004, revealed a new functions of diterpenoids as an anti-proteolytic and anti-haemorrhagic. These authors isolated diterpenoids from the medicinal plant called *Baccharis trimera* and were tested against *Bothrops jararacussu* venom. Another report done by Domingos et al., 2011 talks about the anti-snake activity of diterpenes isolated from marine brown algae (*Carnistrocarpus cervicornis*). The researchers tested the molecule against PLA2 isolated from the snake *Lachesis muta* venom. It showed a great ability to inhibit hemolysis, proteolysis, haemorrhage and coagulation. According to Domingos et al., 2011 the mechanism of action can be explained by that diterpenes bind to the sites of catalytic venom enzymes or by chelating metal ions which are important for their enzymatic activity due to its ability to bind divalent metals.

On the other hand, each of the Frc13 and Frc14 has similar components as that of Frc11. Both fractions were tested individually against the snake venoms, *B. jararaca, E. carinatus, C. atrox* as well as pooled venoms. They both showed parial neutralisation of each of *B. jararaca* and *E. carinatus* venoms. However, they were unable to neutralise neither *C. atrox* nor pooled venoms. Their capacity of neutralising the snake venoms can be explained to the types of Phenolics they have. They both contain Anthocyanins (Cyanidin and Malvidin). They also consist of hydrolysable tannins: Gallic acid. Bark of *P. dulce* has been reported to contain 37% of condensed tannins (Pithayanukul et al., 2005). tannins mechanism and their action are summarized in three points: (i) scavenging free radicals, (ii) chelating trace metals and (iii) binding protein and inhibit their enzymatic activity. The antioxidant property of tannins is similar to that of flavonoids. They prevent cardiovascular diseases by several mechanisms: antioxidative, antithrombogenic and anti-inflammatory (Pereira et al., 2009). Sia and colleagues 2011, isolated tannins from *Mimosa pudica* to test against cobra (*Naja kaouthia*) venom. They found out that the preincubation of both tannins and snake venom result in a protection of mice from death. However when tannins was introduced after injecting the same snake venom, it failed to prevent mice death (Sia et al., 2011). The same thing was reported by Moura et al., 2014, that the preincubation of tannins with *B. jararaca* venom inhibits the haemorrhagic activity of the venom. However, when the tannins has been introduced orally, it showed no activity against the venom which has been introduced intradermally. The authors explained that to the nature of tannins itself. Tannins cannot be hydrolysed due to the polymers of flavan-3-ols (catechin) and/or flavan 3,4 diols (leucoanthocyanidin) linked by carbon– carbon bonds, therefore it cannot be absorbed by gastrointestinal tract. Hence the preincubation allow tannins to chelate metal ions which are essential for some snake venom enzymes e.g. zinc SVMP (Moura et al., 2014) and precipitating them (Silva et al., 2014). Flavonoids demonstrate as well metal chelating activity by promoting strong hydrogen bonds with amides of protein chains (Silva et al., 2014).

When the three active fractions were combined together, they showed a complete neutralisation activity. They were able to neutralise *C. atrox* as well as the pooled venoms, which they were not be able to neutralise individually. The activity can be explained by that, increasing of the polyphenols and hydrolysable tannins concentration allow better neutralising activity in combination with the Acyclic diterpenoids which present in Frc11. Those secondary metabolites have the ability to cross react and bind with enzymatic components of the venom as well as inhibiting the other protein contents found in venom (Kumarapppan et al., 2011).

The anti-lethal activity of the active fractions was performed using LD_100_. An amount of 30μl (0.026mg/mL), 80μl (0.021mg/mL) and 80μl (0.018mg/mL) were taken from Frc11, Frc13, and Frc14, respectively. They were preincubated with LD_100_ and introduced i.p. The anti-lathality of the active fractions was statistically not significant. The negative control which received venom only died after 18 mintues. While those which received crude extract (20mg/kg) survived for 35 minutes. Group of mice which received Frc11 survived for at most 24 minutes. While the group which received Frc13 survived for about 37 minutes. The last group which received Frc14 survived for at most 33 minutes. The group which received the commercial Saudi antivenom died at 29 minutes later. While the group of the commercial Indian antivenom, survived the longest period for at most 48 minutes.

The elongation of the survival time of mice which received the active fraction, especially Frc13, were noticed to be more comparing with those that received the crude extract. And that’s due to the elimination of toxic components via fractionation. The pooled fractions were not tested against LD_100_ due to the limitation of the samples. However, my prediction is that, if I have enough samples then I believed with bring promising results and masking the positive controls used, where the survival rate would confirm this study postulated hypothesis.

Fraction samples were also examined their neutralisation efficacy to inhibit the snake venom gelatinase activity using gelatin zymography following the protocol of Hasson et al., 2003. The gel was incorporated with gelatin and the different fractions were preincubated with *B. jararaca* venom to evaluate their anti gelatinase activity. After electrophoresis, the individual fractions did not neutralise snake venom. However when the same fractions were combined, they showed an inhibition of gelatinase enzyme.

## References

1. Arora, V., & Choudhary, C. (2016). Cortical Blindness After Snake Bite Envenomation. International Journal of Scientific Research, 5(4).

2. Barreto, G. N. L. S., de Oliveira, S. S., dos Anjos, I. V., de Menezes Chalkidis, H., Mourão, R. H. V., da Silva, A. M. M., … de Camargo Gonçalves, L. R. (2017). Experimental Bothrops atrox envenomation: Efficacy of antivenom therapy and the combination of Bothrops antivenom with dexamethasone. PLoS Neglected Tropical Diseases, 11(3), e0005458.

3. Bordon, K. C., Wiezel, G. A., Cabral, H., & Arantes, E. C. (2015). Bordonein-L, a new L-amino acid oxidase from Crotalus durissus terrificus snake venom: isolation, preliminary characterization and enzyme stability. Journal of Venomous Animals and Toxins including Tropical Diseases, 21(1), 26.

4. Bhattacharjee, P., & Bhattacharyya, D. (2013). Medicinal plants as snake venom antidotes. J Exp Appl Anim Sci, 1, 156–181.

5. Calvete, J. J., Sanz, L., Angulo, Y., Lomonte, B., & Gutiérrez, J. M. (2009). Venoms, venomics, antivenomics. FEBS letters, 583(11), 1736–1743.

6. Cardoso, J. L. C., Fan, H. W., França, F. O., Jorge, M. T., Leite, R. P., Nishioka, S. A., … & Chudzinski, A. M. (1993). Randomized comparative trial of three antivenoms in the treatment of envenoming by lance-headed vipers (Bothrops jararaca) in São Paulo, Brazil. QJM: An International Journal of Medicine, 86(5), 315–325

7. Chippaux, J.-P. (1998). The development and use of immunotherapy in Africa. Toxicon, 36(11), 1503–1506.

8. Chippaux, J.-P. (2008). Estimating the global burden of snakebite can help to improve management. PLoS Med, 5(11), e221.

9. Domingos, T. F. S., Vallim, M. A., Carvalho, C., Sanchez, E. F., Teixeira, V. L., & Fuly, A. L. (2011). Anti-snake venom effect of secodolastane diterpenes isolated from Brazilian marine brown alga Canistrocarpus cervicornis against Lachesis muta venom. Revista Brasileira de Farmacognosia, 21(2), 234–238.

10. Gopi, K., Renu, K., Vishwanath, B. S., & Jayaraman, G. (2015). Protective effect of Euphorbia hirta and its components against snake venom induced lethality. Journal of ethnopharmacology, 165, 180–190.

11. Gopi, K., Anbarasu, K., Renu, K., Jayanthi, S., Vishwanath, B. S., & Jayaraman, G. (2016). Quercetin-3-O-rhamnoside from Euphorbia hirta protects against snake Venom induced toxicity. Biochimica et Biophysica Acta (BBA)-General Subjects, 1860(7), 1528–1540.

12. Gupta, Y., & Peshin, S. (2012). Do herbal medicines have potential for managing snake bite envenomation? Toxicology international, 19(2), 89.

13. Gutiérrez, J. M., Theakston, R. D. G., & Warrell, D. A. (2006). Confronting the neglected problem of snake bite envenoming: the need for a global partnership. PLoS Med, 3(6), e150.

14. Gutiérrez, J. M., León, G., & Lomonte, B. (2003). Pharmacokinetic-pharmacodynamic relationships of immunoglobulin therapy for envenomation. Clinical pharmacokinetics, 42(8), 721–741.

15. Gutiérrez, J. M., León, G., & Burnouf, T. (2011). Antivenoms for the treatment of snakebite envenomings: the road ahead. Biologicals, 39(3), 129–142.

16. Hasson, S., Theakston, R. D. G., & Harrison, R. (2004). Antibody zymography: a novel adaptation of zymography to determine the protease-neutralising potential of specific antibodies and snake antivenoms. Journal of immunological methods, 292(1), 131–139.

17. Hifumi, T., Sakai, A., Kondo, Y., Yamamoto, A., Morine, N., Ato, M., … Kato, H. (2015). Venomous snake bites: clinical diagnosis and treatment. Journal of intensive care, 3(1), 16.

18. Hossain, M. A., AL-Raqmi, K. A. S., AL-Mijizy, Z. H., Weli, A. M., & Al-Riyami, Q. (2013). Study of total phenol, flavonoids contents and phytochemical screening of various leaves crude extracts of locally grown Thymus vulgaris. Asian pacific journal of tropical biomedicine, 3(9), 705–710.

19. Isbister, G. K., Brown, S. G., MacDonald, E., White, J., & Currie, B. J. (2008). Current use of Australian snake antivenoms and frequency of immediate-type hypersensitivity reactions and anaphylaxis. Medical Journal of Australia, 188(8), 473.

20. Ismail, M., Abd-Elsalam, M. A., & Al-Ahaidib, M. S. (1998). Pharmacokinetics of 125 I-labelled Walterinnesia aegyptia venom and its specific antivenins: flash absorption and distribution of the venom and its toxin versus slow absorption and distribution of IgG, F (ab’) 2 and F (ab) of the antivenin. Toxicon, 36(1), 93–114.

21. Janardhan, B., Shrikanth, V. M., Mirajkar, K. K., & More, S. S. (2014). In vitro screening and evaluation of antivenom phytochemicals from Azima tetracantha Lam. leaves against Bungarus caeruleus and Vipera russelli. Journal of Venomous Animals and Toxins including Tropical Diseases, 20(1), 12.

22. Januário, A. H., Santos, S. L., Marcussi, S., Mazzi, M. V., Pietro, R. C., Sato, D. N., … & Soares, A. M. (2004). Neo-clerodane diterpenoid, a new metalloprotease snake venom inhibitor from Baccharis trimera (Asteraceae): anti-proteolytic and anti-hemorrhagic properties. Chemico-biological interactions, 150(3), 243–251.

23. Kalavani, R., Banu, R. S., Jeyanthi, K., Sankari, T. U., & Kanna, A. V. (2016). Evaluation of anti-inflammatory and antibacterial activity of Pithecellobium dulce (Benth) extract. Biotechnological Research, 2(4), 148–154.

23. Kuruppu, S., Smith, A. I., Isbister, G. K., & Hodgson, W. C. (2008). Neurotoxins from Australo-Papuan elapids: a biochemical and pharmacological perspective. Critical reviews in toxicology, 38(1), 73–86.

24. Kasturiratne, A., Wickremasinghe, A. R., de Silva, N., Gunawardena, N. K., Pathmeswaran, A., Premaratna, R., … de Silva, H. J. (2008). The global burden of snakebite: a literature analysis and modelling based on regional estimates of envenoming and deaths. PLoS Med, 5(11), e218.

25. Khanzada, S. K., Khanzada, A. K., Shaikh, W., & Ali, S. A. (2013). Phytochemical studies on Pithecellobium dulce benth. a medicinal plant of Sindh, Pakistan. Pak. J. Bot, 45(2), 557–561.

26. Kumarapppan, C., Jaswanth, A., & Kumarasunderi, K. (2011). Antihaemolytic and snake venom neutralizing effect of some Indian medicinal plants. Asian Pacific journal of tropical medicine, 4(9), 743–747.

27. León, G., Valverde, J. M., Rojas, G., Lomonte, B., & Gutiérrez, J. M. a. (2000). Comparative study on the ability of IgG and Fab sheep antivenoms to neutralize local hemorrhage, edema and myonecrosis induced by Bothrops asper (terciopelo) snake venom. Toxicon, 38(2), 233–244.

28. Lim, T. (2012). Quercus infectoria *Edible Medicinal And Non-Medicinal Plants* (pp. 16–26): Springer

29. Lipps, B. V., & Lipps, F. W. (1996). U.S. Patent No. 5,576,297. Washington, DC: U.S. Patent and Trademark Office.

30. Maduwage, K., Silva, A., O’Leary, M. A., Hodgson, W. C., & Isbister, G. K. (2016). Efficacy of Indian polyvalent snake antivenoms against Sri Lankan snake venoms: lethality studies or clinically focussed in vitro studies. Scientific reports, 6.

31. Manna, P., Bhattacharyya, S., Das, J., Ghosh, J., & Sil, P. C. (2011). Phytomedicinal role of Pithecellobium dulce against CCl4-mediated hepatic oxidative impairments and necrotic cell death. Evidence-based complementary and alternative medicine, 2011.

32. Megala, J., & Geetha, A. (2010). Free radical-scavenging and H+, K+-ATPase inhibition activities of Pithecellobium dulce. Food chemistry, 121(4), 1120–1128.

33. Menaldo, D. L., Jacob-Ferreira, A. L., Bernardes, C. P., Cintra, A. C., & Sampaio, S. V. (2015). Purification procedure for the isolation of a PI metalloprotease and an acidic phospholipase A 2 from Bothrops atrox snake venom. Journal of Venomous Animals and Toxins including Tropical Diseases, 21(1), 28.

34. Mendes, M. M., Vieira, S. A. P. B., Gomes, M. S. R., Paula, V. F., Alcântara, T. M., Homsi-Brandeburgo, M. I., … & Rodrigues, V. M. (2013). Triacontyl p-coumarate: An inhibitor of snake venom metalloproteinases. Phytochemistry, 86, 72–82.

35. Moreira, V., Teixeira, C., da Silva, H. B., Lima, M. R. D. I., & Dos-Santos, M. C. (2016). The role of TLR2 in the acute inflammatory response induced by Bothrops atrox snake venom. Toxicon, 118, 121–128.

36. Moura de, V. M., Bezerra, A. N. S., Mourão, R. H. V., Lameiras, J. L. V., Raposo, J. D. A., de Sousa, R. L., … & Dos-Santos, M. C. (2014). A comparison of the ability of Bellucia dichotoma Cogn.(Melastomataceae) extract to inhibit the local effects of Bothrops atrox venom when pre-incubated and when used according to traditional methods. Toxicon, 85, 59–68.

37. Nagmoti, D. M., Kothavade, P. S., Bulani, V. D., Gawali, N. B., & Juvekar, A. R. (2015). Antidiabetic and antihyperlipidemic activity of Pithecellobium dulce (Roxb.) Benth seeds extract in streptozotocin-induced diabetic rats. European Journal of Integrative Medicine, 7(3), 263–273.

38. Owuor, B. O., & Kisangau, D. P. (2006). Kenyan medicinal plants used as antivenin: a comparison of plant usage. Journal of ethnobiology and ethnomedicine, 2(1), 7.

39. Pereira, D. M., Valentão, P., Pereira, J. A., & Andrade, P. B. (2009). Phenolics: From chemistry to biology.

40. Pithayanukul, P., Ruenraroengsak, P., Bavovada, R., Pakmanee, N., Suttisri, R., & Saen-oon, S. (2005). Inhibition of Naja kaouthia venom activities by plant polyphenols. Journal of ethnopharmacology, 97(3), 527–533.

41. Preethi, S., & Saral, M. (2016). Screening of natural polysaccharides extracted from the fruits of Pithecellobium dulce as a pharmaceutical adjuvant. International Journal of Biological Macromolecules, 92, 347–356.

42. Samy, R. P., Gopalakrishnakone, P., & Chow, V. T. (2012). Therapeutic application of natural inhibitors against snake venom phospholipase A2. Bioinformation, 8(1), 48.

43. Shabbir, A., Shahzad, M., Masci, P., & Gobe, G. C. (2014). Protective activity of medicinal plants and their isolated compounds against the toxic effects from the venom of Naja (cobra) species. Journal of ethnopharmacology, 157, 222–227.

44. Shenoy, P. A., Nipate, S. S., Sonpetkar, J. M., Salvi, N. C., Waghmare, A. B., & Chaudhari, P. D. (2014). Production of high titre antibody response against Russell’s viper venom in mice immunized with ethanolic extract of fruits of Piper longum L.(Piperaceae) and piperine. Phytomedicine, 21(2), 159–163.

45. Sia, F. Y., Vejayan, J., Jamuna, A., & Ambu, S. (2011). Efficacy of tannins from Mimosa pudica and tannic acid in neutralizing cobra (Naja kaouthia) venom. Journal of Venomous Animals and Toxins including Tropical Diseases, 17(1), 42–48.

46. Silva-Félix, J., Souza, T., Menezes, Y. A., Cabral, B., Câmara, R. B., Silva-Junior, A. A., … & Fernandes-Pedrosa, M. F. (2014). Aqueous leaf extract of Jatropha gossypiifolia L.(Euphorbiaceae) inhibits enzymatic and biological actions of Bothrops jararaca snake venom. PLoS One, 9(8), e104952.

47. Singhal, N., Kumar, M., Kanaujia, P. K., & Virdi, J. S. (2015). MALDI-TOF mass spectrometry: an emerging technology for microbial identification and diagnosis. Frontiers in microbiology, 6.

48. Stock, R. P., Massougbodji, A., Alagón, A., & Chippaux, J.-P. (2007). Bringing antivenoms to sub-Saharan Africa. Nature biotechnology, 25(2), 173–177.

49. Sugumaran, M., Vetrichelvan, T., & Quine, S. D. (2008). Locomotor Activity of Leaf Extracts of Pithecellobium dulce Benth. Ethnobotanical Leaflets, 2008(1), 62.

50. WHO, W. H. O. (2007). Rabies and envenomings: a neglected public health issue. Report of a consultative meeting, World Health Organization, Geneva, 10 January 2007. Paper presented at the Rabies and envenomings: a neglected public health issue. Report of a consultative meeting, World Health Organization, Geneva, 10 January 2007.

51. Tanwar, P., Ghorui, S., Kochar, S., Singh, R., & Patil, N. (2017). Production and preclinical assessment of camelid immunoglobulins against Echis sochureki venom from desert of Rajasthan, India. Toxicon, 134, 1–5.

52. Tang, X. N., Berman, A. E., Swanson, R. A., & Yenari, M. A. (2010). Digitally quantifying cerebral hemorrhage using Photoshop® and Image J. Journal of neuroscience methods, 190(2), 240–243.

53. Theakston, R. D. G. (1997). 18 The kinetics of snakebite envenoming and therapy. In Symposia of the Zoological Society of London (No. 70, pp. 251-260). London: The Society, 1960–1999.

