## Supplement data for "Composition efficacy of Unsaturated Arachidonic acid, Diterpenoids, Malvin (C_29_H_35_ClO_17_), and Bergenin to neutralise venom from different venomous snake species"

*Keywords: Pithecellobium dulce; B. jararaca: Bothrops jararaca; C. atrox: Crotalus atrox; E. carinatus: Echis carinatus; E/e: crude exthanolic extract; DF: dilution factor; s.c.: subcutaneously; i.p.: intaperitoneally;* $LD_{50}$*: median lethal dose; HPLC: high performance liquid chromatography; Frc: fraction; SVMP: snake venom metalloproteinase; MALDI-TOF MS: matrix-assisted laser desorption/ionization-time of flight mass spectrometer;* $LD_{100}$*: complete lethal dose; IAV: Indian antivenom.*

### Other Supplemented Results from Results Analysis

### RESULTS& ANALYSIS

#### Table 4. 1: The number and distribution of mice used in the experiments

| Types of experiments | Testing mice n= 211 | Control mice n= 82 |
|---|---|---|
| **MHD** | **6** | **-** |
| **Screening of extracts** | **18** | **2** |
| **Acute toxicity test** | **40** | **10** |
| **Determination of the ED of E/e** | **8** | **2** |
| **Screening of fractions** | **63** | **8** |
| **Evaluating the activity of the powerful fractions** | **36** | **36** |
| **Anti-lethal activity** | **40** | **24** |
| ***MHD: Minimum haemorrhagic dose; ED: Effective dose; E/e: Ethanolic/ethanolic extract*** | | |

#### Table 4. 2: Differentiation criteria between mild, to severe hemorrhage

| Range | Description | Sign |
|---|---|---|
| **Area ≤ 0.6 mm$^2$** | No haemorrhage | - |
| **0.6 > Area ≤ 30 mm$^2$** | Mild haemorrhage | + |
| **30>Area ≤ 50 mm$^2$** | Heamorrhage | ++ |
| **Area > 50 mm$^2$** | Severe haemorrhage | +++ |

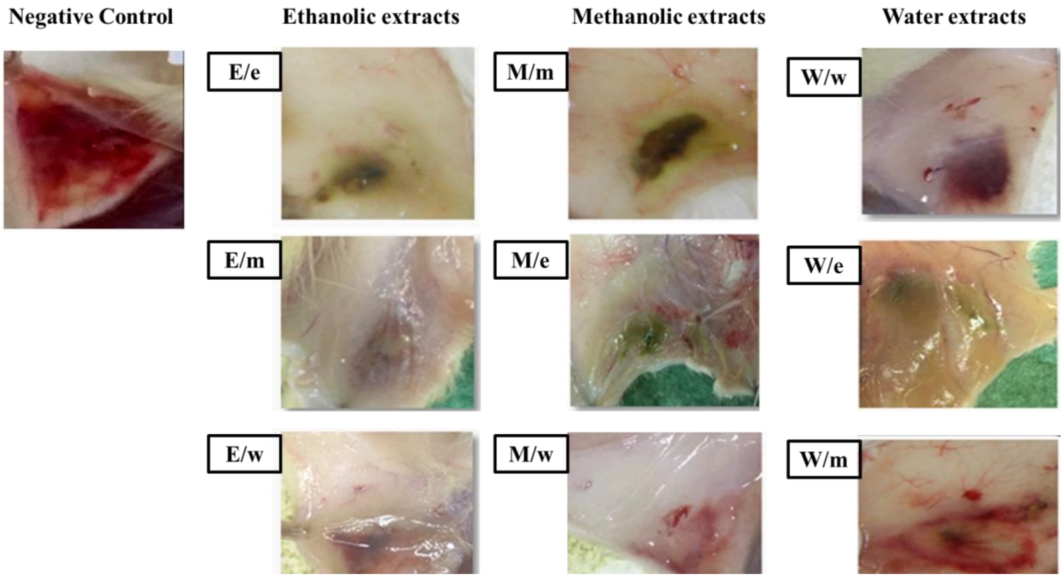


**Figure 4. 1: Verification of the active extract.** *E/e: Ethanolic/ethanolic; E/m: Ethanolic/methanolic; E/w: Ethanolic/water; M/m: Methanolic/methanolic; M/e: Methanolic/ethanolic; M/w: Methanolic/water; W/w: Water/water; W/e: Water/ethanolic; W/m: Water/methanolic.*

**Table 4. 3: The severity of haemorrhage of the different solvent extracts in Balab/C mice**

| Type of sample | Rate of hemorrhage Severity ($mm^2$) | |
|---|---|---|
| | Group 1 | Group 2 |
| E/e | - | - |
| E/m | + | + |
| E/w | + | + |
| M/m | - | ++ |
| M/e | + | + |
| M/w | + | + |
| W/w | +++ | +++ |
| W/e | + | + |
| W/m | +++ | +++ |

*E/e: Ethanolic/ethanolic; E/m: Ethanolic/methanolic; E/w: Ethanolic/water; M/m: Methanolic/methanolic; M/e: Methanolic/ethanolic; M/w: Methanolic/water; W/w: Water/water; W/e: Water/ethanolic; W/m: Water/methanolic.*

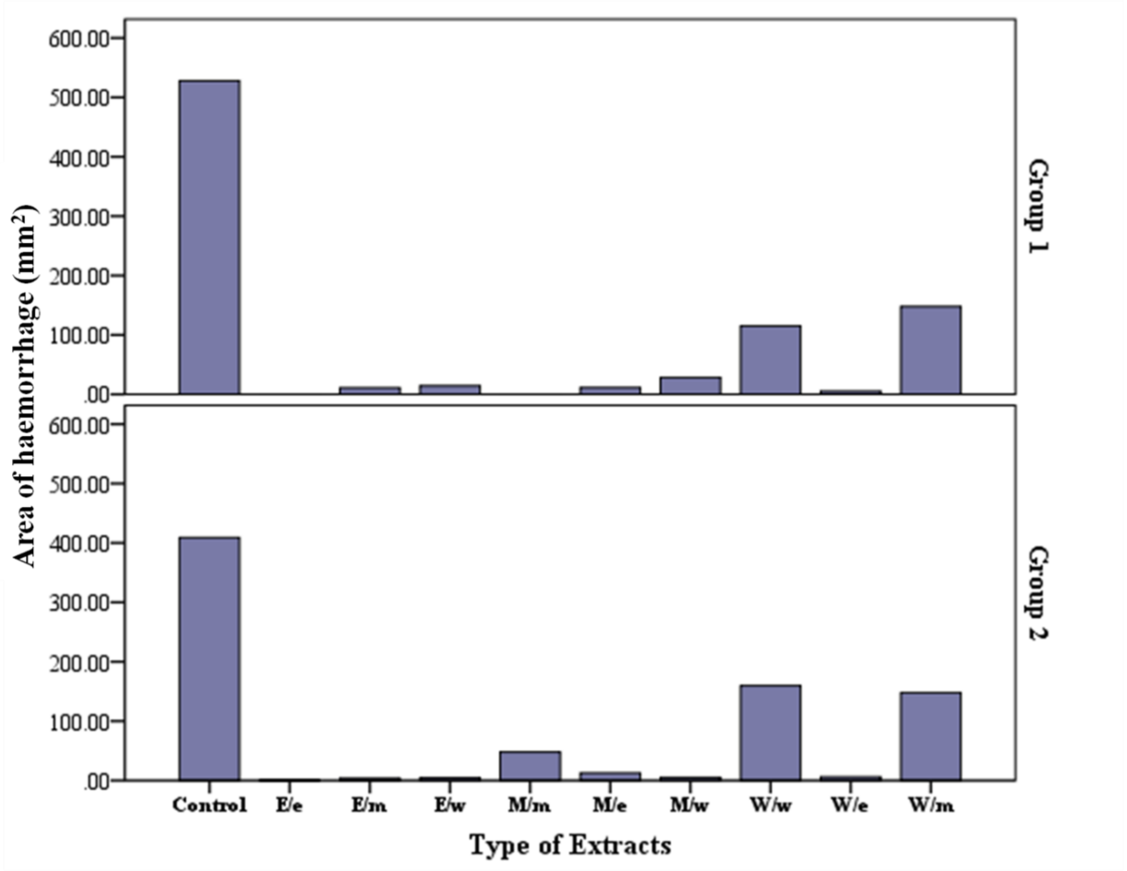


4.5

**Figure 4. 2: Screening of different solvent extracts.** *E/e: Ethanolic/ethanolic; E/m: Ethanolic/methanolic; E/w: Ethanolic/water; M/m: Methanolic/methanolic; M/e: Methanolic/ethanolic; M/w: Methanolic/water; W/w: Water/water; W/e: Water/ethanolic; W/m: Water/methanolic.*

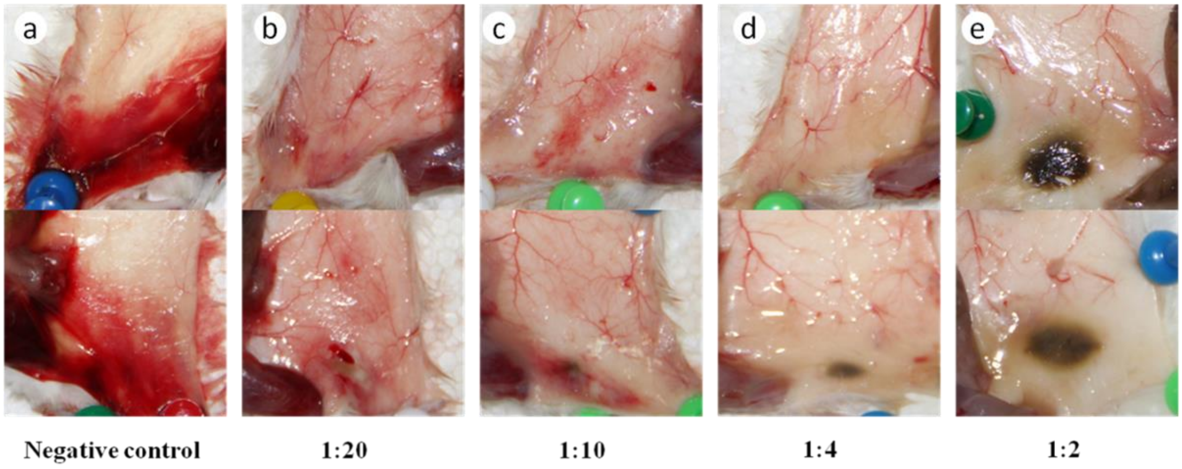


**Figure 4. 3: Histological sections show efficacy of different DFs of the E/e crude extract to neutralise the local haemorrhage induced by *B. jararaca*.** a) represents the negative control which contains venom only. While, (b-e) are the preincubated mixture of different concentrations of E/e with the venom.

**Table 4. 4: Severity of haemorrhage of the DFs of E/e**

| DF of E/e | Severity. Haem | |
|---|---|---|
| | Group 1 | Group 2 |
| **1:20** | + | ++ |
| **1:10** | + | + |
| **1:4** | - | - |
| **1:2** | - | - |
| *DF: dilution factor; E/e: ethanolic/ethanolic* | | |

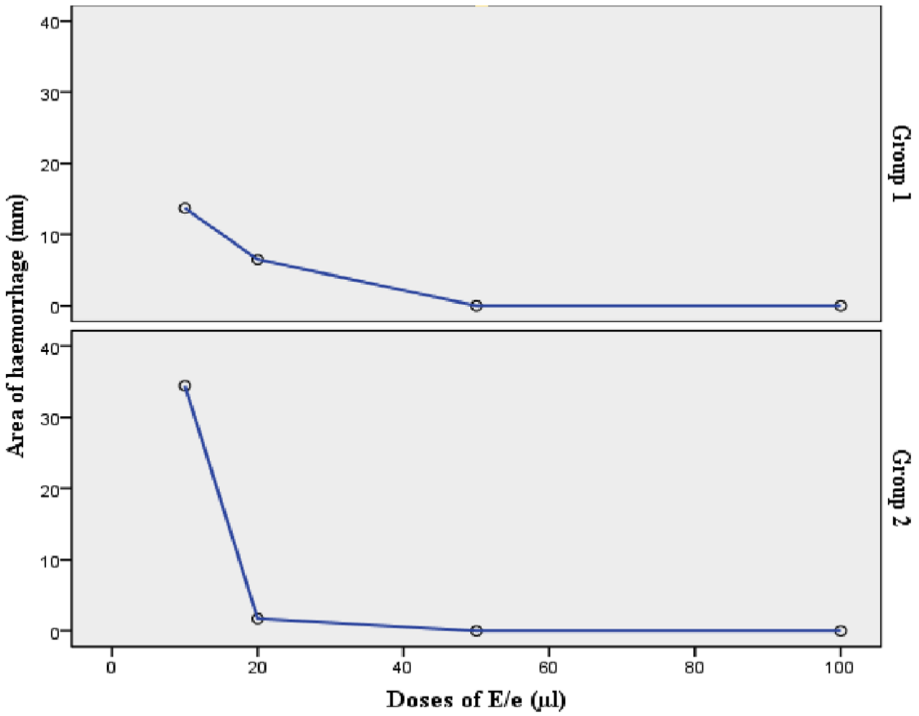


**Figure 4. 4: Linear relationship of the different DFs of E/e with their neutralisation efficacy**

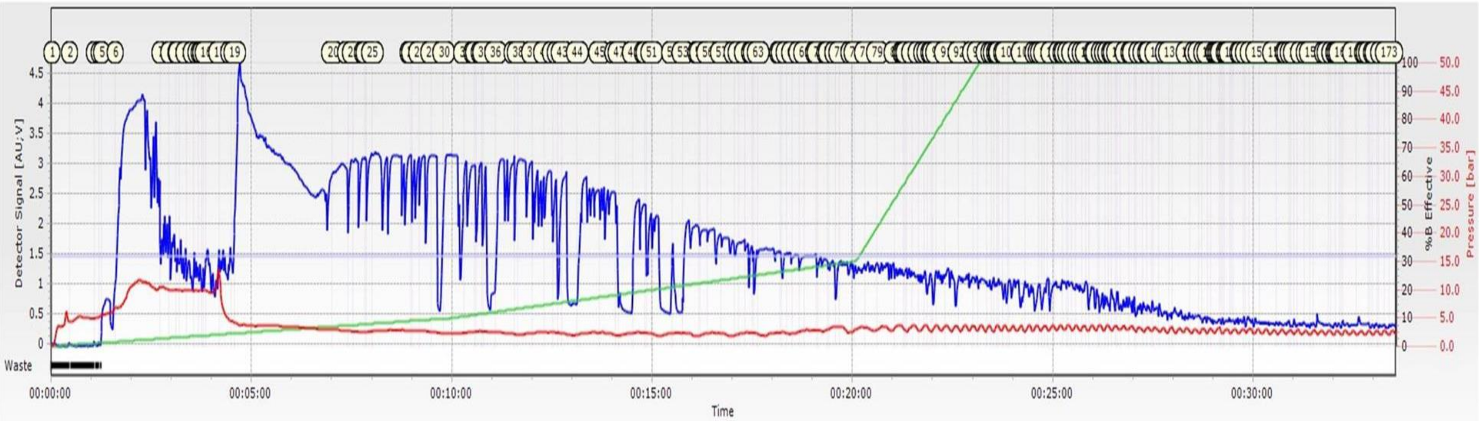


**Figure 4. 5: The fractions obtained from HPLC chromatography.** The graph shows the 173 fractions obtained. The blue peaks represent the absorbance of each fraction at 280nm, which was used to obtain the concentration of the active fractions.

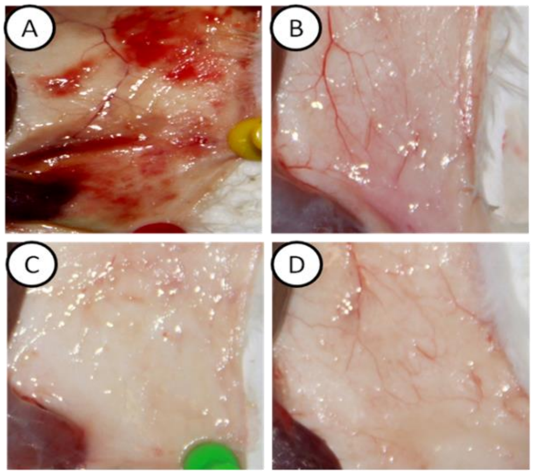


**Figure 4. 6: Evaluation of groups A, D and H against *B. jararaca*.** (A) negative control, (B) group A, (C) group D and (D) group H.

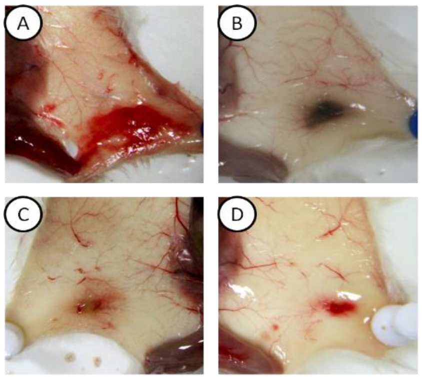


**Figure 4. 7: Evaluation of groups A, D and H against pooled venoms.** (A) negative control, (B) group A, (C) group D and (D) group H.

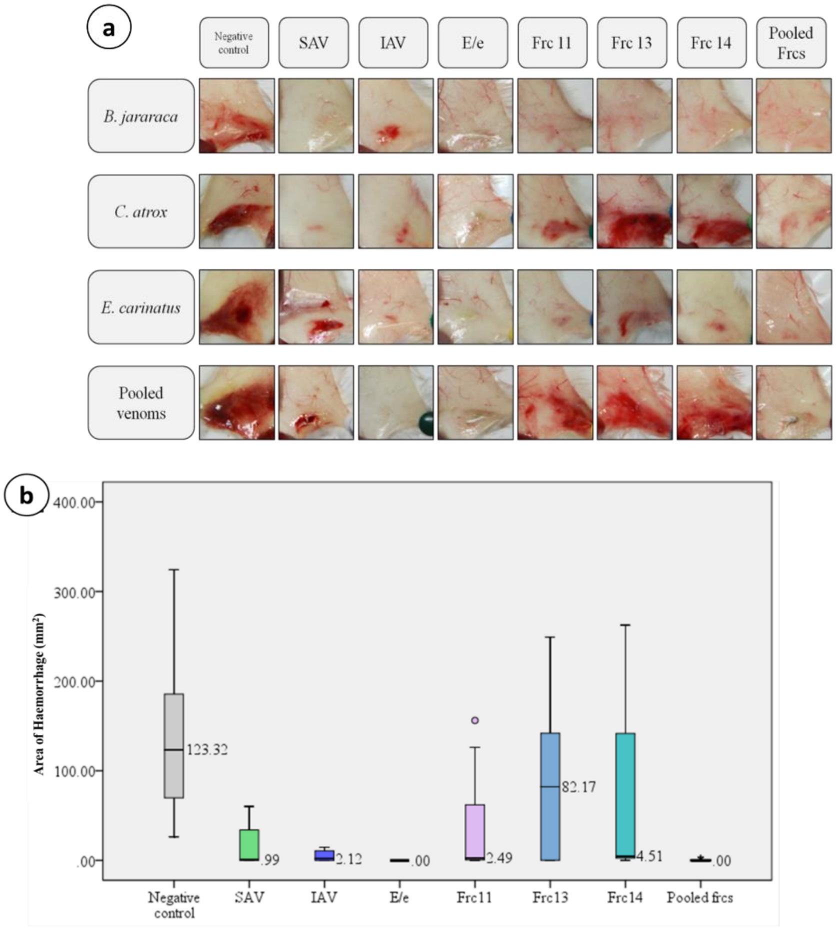


**Figure 4. 8: evaluation the efficacy of the active fractions 11, 13 and 14 against B. jararaca, C. atrox, E. carinatus as well as pooled venoms.** (a) Shows the histological samples mice dissected skins. (b) neutralisation activity of the fractions 11, 13, 14 as well as the pooled fractions to neutralise different venoms in comparison with the positive and negative controls. Box plot shows the area of haemorrhage. The graph clearly shows that fraction 11 has the best neutralisation activity compared with the other two fractions. When the fractions are pooled together, they showed a complete netralisation of the venoms. *SAV: Saudi antivenom; IAV: Indian antivenom; E/e: Ethanolic/ethanolic; Frc/s: Fraction/s.*

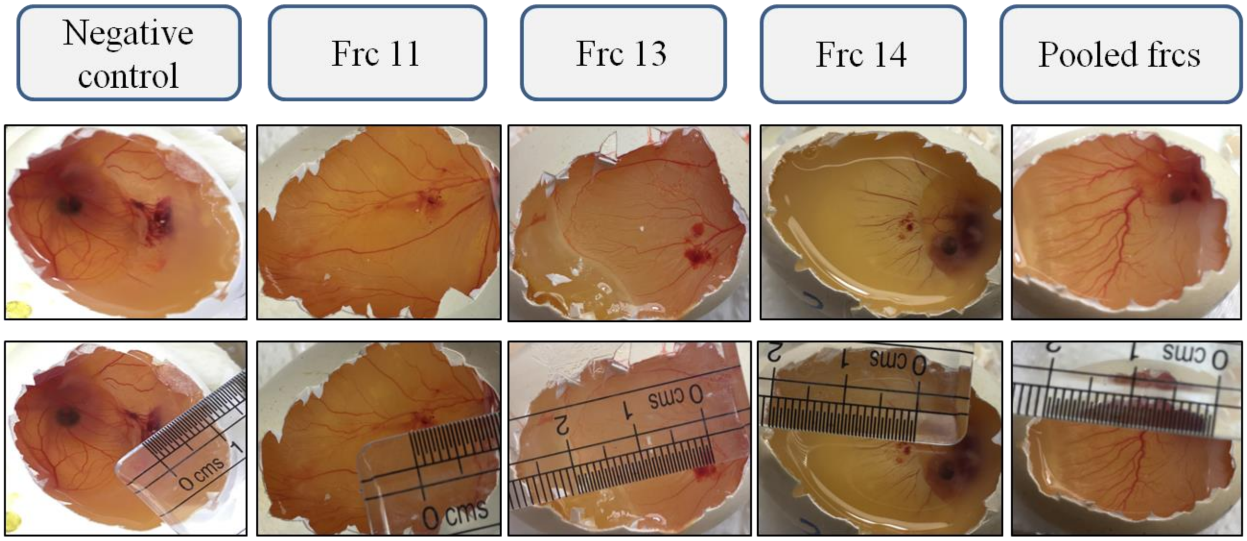


**Figure 4. 9: Evaluation of the active fractions using egg embryo.** Frc11 showed the best neutralisation as an individual fraction in comparison with the other two fractions. Pooled fractions have the greatest activity.

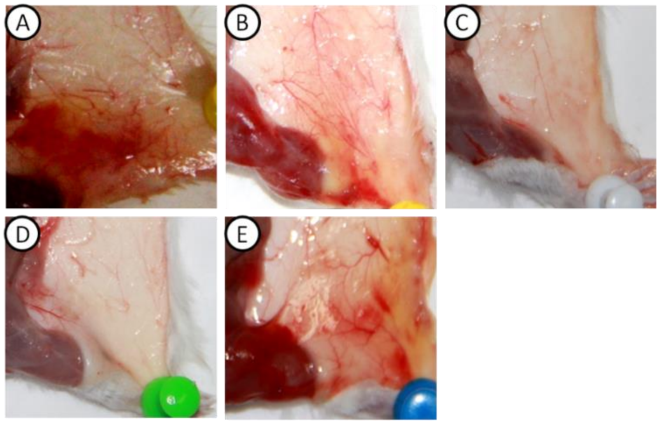


**Figure 4. 10: Evaluation the activity of the subgroups A1a, A1b, A2a and A2b against *B. jararaca* venom.** (A) negative control, (B) subgroup A1a (1-5), (C) subgroup A1b (6-10), (D) subgroup A2a (11-15) and (E) subgroup A2b (16-20).

**Table 4. 5: The different components of the peaks of Frc11**

| S. No. | Database | | Chemical structure |
|---|---|---|---|
| 1 | **Chemical name** | Unsaturated arachaetidic acid | 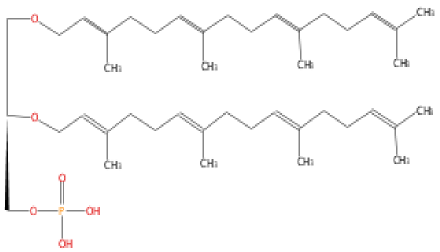 |
| | **Class** | Prenol lipids | |
| | **Direct parent** | Acyclic diterpenoids | |
| | **Molecular formula** | $C_{43}H_{73}O_6P$ | |
| | **Total exact mass** | 716.5145 | |

**Table 4. 6: The components of the peaks presented by both Frc13 and Frc14**

| S. No. | Database | | Chemical structure |
|---|---|---|---|
| 1 | **Chemical name** | Malvin | 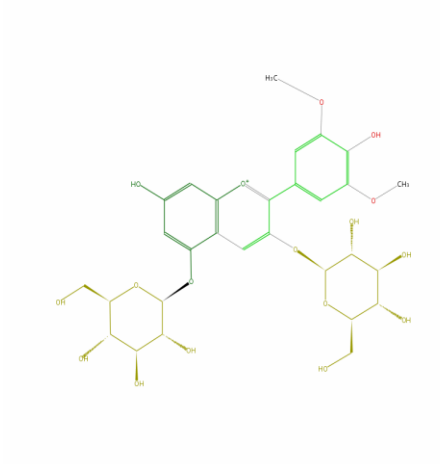 |
| | **Class** | Flavonoids | |
| | **Direct parent** | Anthocyanidin-5-O-glycosides | |
| | **Molecular formula** | $C_{29}H_{35}O_{17}$ | |
| | **Total exact mass** | 655.1869 | |
| 2 | **Chemical name** | Bergenin | |
| | **Class** | Hydrolysable tannin | |
| | **Direct parent** | Gallic acid | |

| | | | |
|---|---|---|---|
| | **Molecular formula** | $C_{14}H_{16}O_{9}$ | 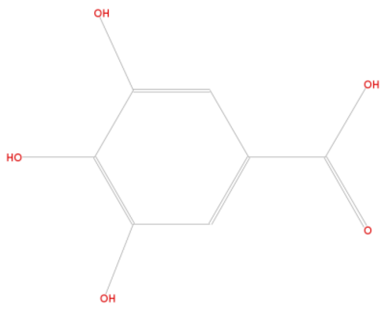 |
| | **Total exact mass** | 328.0794 | |
| **3** | **Chemical name** | Cy 3-coumSamb-5-Glc | 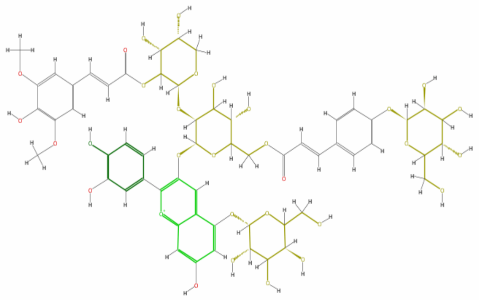 |
| | **Class** | Flavonoids | |
| | **Direct parent** | Anthocyanidin 3-O-6-p-coumaroyl glycoside | |
| | **Molecular formula** | $C_{41}H_{45}O_{22}$ | |
| | **Total exact mass** | 889.2397 | |

**Table 4. 7: The anti-median lethal activity of the different concentrations of E/e**

| Type. Sample | *B. jararaca* (1.1 mg/ kg) Venom conc. (5 µg/ 1 µl) Dose per mouse (µg) | Doses/mouse ( µl) | Survival/No. mice | Survival % | *p* value ($p$<0.05) |
|---|---|---|---|---|---|
| **-ve Control** | 27.50 | - | 0/4 | 0% | |
| **SAV** | 27.50 | 193.4 | 4/4 | 100% | 0.008 |
| **IAV** | 27.50 | 193.4 | 4/4 | 100% | 0.008 |
| **E/e** | 27.50 | 10 | 3/4 | 75% | 0.040 |
| **E/e** | 27.50 | 20 | 2/4 | 50% | 0.127 |
| **E/e** | 27.50 | 40 | 3/4 | 75% | 0.040 |
| **E/e** | 27.50 | 60 | 2/4 | 50% | 0.127 |
| **E/e** | 27.50 | 100 | 1/4 | 25% | 0.317 |
| **E/e** | 27.50 | 180 | 1/4 | 25% | 0.317 |

*SAV: Saudi antivenom; IAV: Indian antivenom; E/e: Ethanolic/ethanolic extract.*

**Table 4. 8: The anti-total lethal activity of the E/e and the active fractions**

| Type. Sample | *Pooled venom* *Bj+Ca+Ec* Dose (µg/ mouse) | Doses/mouse | No. of mice | Survival/mice | Survival % | Average survival time (minute) |
|---|---|---|---|---|---|---|
| **Control** | 180 | - | 4 | 0/4 | 0% | 18 |
| **SAV** | 180 | 193.4 | 4 | 0/4 | 0% | 29 |
| **IAV** | 180 | 193.4 | 4 | 0/4 | 0% | 48 |
| **E/e** | 180 | 80 | 4 | 0/4 | 0% | 35 |
| **Frc11** | 180 | 30 | 4 | 0/4 | 0% | 24 |
| **Frc13** | 180 | 80 | 4 | 0/4 | 0% | 37 |
| **Frc14** | 180 | 80 | 4 | 0/4 | 0% | 33 |

*SAV: Saudi antivenom; IAV: Indian antivenom; E/e: Ethanolic/ethanolic extract.*
